# *Bacteroides ovatus* administration enhanced α-mPD-1 immunotherapy in a mouse colorectal cancer model

**DOI:** 10.64898/2026.11.24.769059

**Authors:** Huan Zhang, Shiqi Zhang, Heng-Yi Chen, Shuaixin Gao, Mengyang Xu, Xinru Pang, Li Chen, Hyein Jung, Chao Guo, Andrew Gold, Natalie Crawford, Fouad Choueiry, Owen Laib, Haifei Shi, Chan-Wang Jerry Lio, Jiangjiang Zhu

**Author notes:** Corresponding author: Jiangjiang Zhu, Ph.D., Comprehensive Cancer Center and Department of Human Sciences, The Ohio State University, 2255 Kenny Road, Columbus, OH 43210. These authors contributed equally to the work (Huan Zhang, Shiqi Zhang).

## Abstract

The gut microbiota modulates responses to immune checkpoint blockade, yet the metabolic remodeling associated with microbiome-based combination therapies remains poorly understood, limiting insights into the host-microbiome interactions underlying therapeutic responses. Here, we investigated whether *Bacteroides ovatus* (BO) enhances the antitumor efficacy of programmed cell death protein 1 (PD-1) blockade in an MC38 syngeneic colorectal cancer mouse model. BO combined with anti-mouse PD-1 (α-mPD-1) produced the strongest antitumor response, reducing average tumor burden by 53% compared with PBS controls. BO-specific qPCR detected higher BO-associated signals in tumors from BO-exposed groups, while exploratory flow cytometry evaluated treatment-associated immune changes. Paired 16S rRNA profiling revealed *Bacteroides* enrichment in BO-treated groups and treatment-associated shifts in microbial community composition. Tumor metabolomics identified alterations in glycan turnover, activated-donor metabolism, oxidized lipids, and fatty-acid trafficking associated with combination treatment. Serum metabolomics revealed treatment-associated metabolic profiles, while fecal metabolomics further characterized intestinal metabolic changes. Integrated cross-omics analysis identified notable associations linking microbial taxa with metabolites across biological compartments. Together, these findings suggest that BO enhances the antitumor activity of PD-1 blockade and is associated with microbial and metabolic remodeling, providing a foundation for investigating host-microbiome interactions underlying combination immunotherapy.

**Graphical Abstract:** 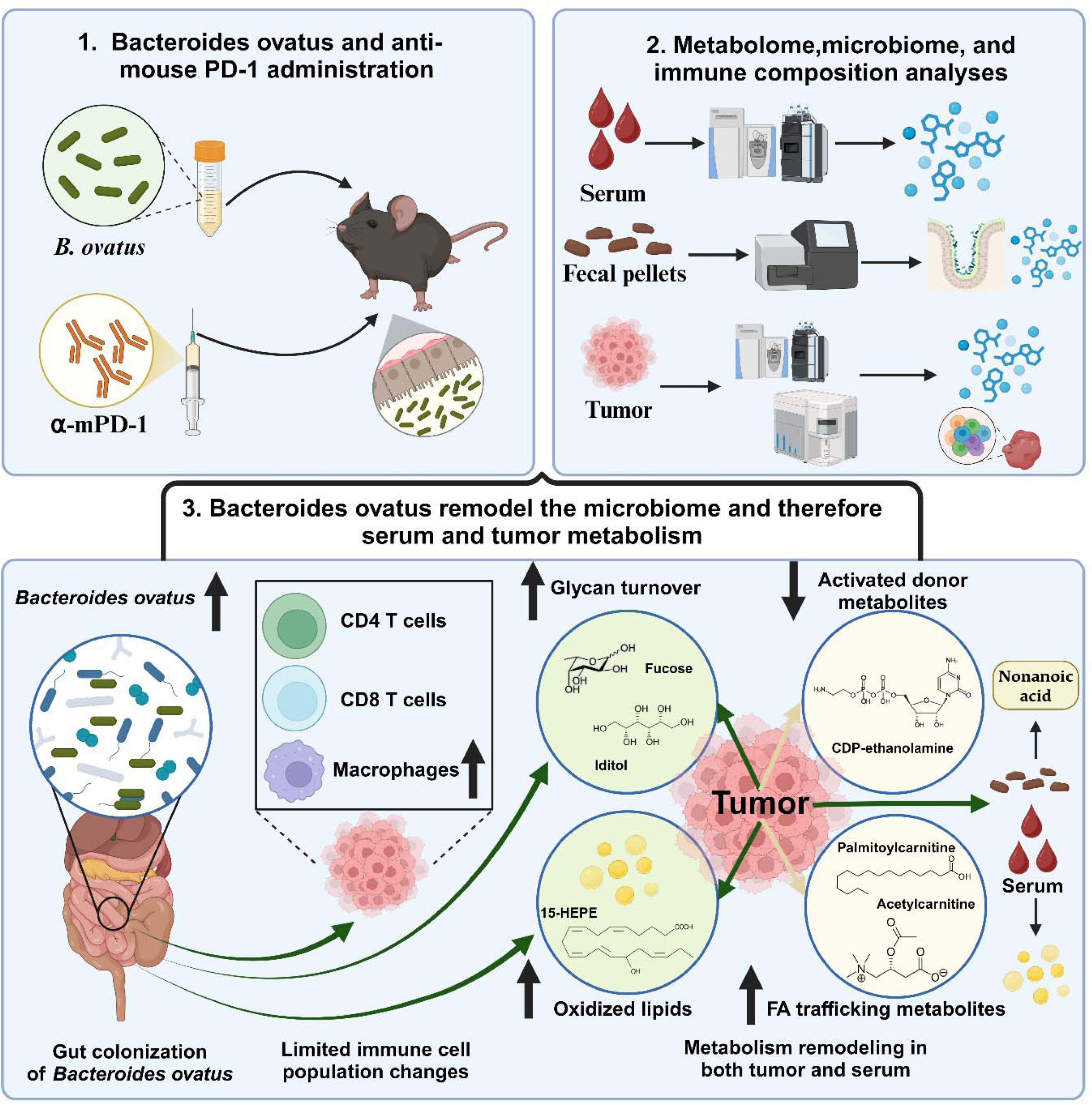

## 1. Introduction

Colorectal cancer (CRC) is a malignancy that arises in the colon or rectum, typically originating from the epithelial cells lining the inner surface of the large intestine.[1] Despite advances in cancer screening, early detection, and surgical intervention, CRC continues to be a significant contributor to global cancer-related mortality.[1] Immune checkpoint blockade targeting programmed cell death protein 1 (PD-1) has changed the cancer treatment landscape for several tumor types and has shown clear benefit in CRC with deficient mismatch repair (MMR) or high microsatellite instability (MSI-H), where pembrolizumab improved progression-free survival compared with chemotherapy in the phase III KEYNOTE-177 study.[2] However, the efficacy of PD-1 blockade in CRC remains inconsistent because most colorectal cancers are MMR proficient and microsatellite stable (MSS), resulting in limited benefit from current checkpoint inhibitor strategies.[3] This therapeutic gap has driven growing interest in identifying host and tumor-extrinsic factors that shape immunotherapy responsiveness in CRC.

Among these factors, the gut microbiome has emerged as a critical player of antitumor immunity.[4,5] Previous research in melanoma and other solid tumors have shown that gut microbial composition is associated with clinical response to anti-PD-1 therapy, and that microbiome transplantation from responders can enhance antitumor effects in preclinical models.[4,6] Subsequent work extended this concept to epithelial tumors and gastrointestinal cancers, supporting the view that microbiome structure is not merely correlative but can be functionally linked to immune checkpoint efficacy.[5,7] In CRC specifically, this question is particularly compelling because the intestinal microbiota is positioned at the interface of mucosal immunity, epithelial biology, and tumor development, and accumulating evidence indicates that gut microbiota dysbiosis contributes to CRC initiation, progression, and therapeutic response.[5,8] Within the CRC field, the role of individual gut bacterial taxa remains complex and context-dependent. Members of the genus *Bacteroides* have been implicated in both beneficial and detrimental host effects depending on species, strain, ecological context, and disease setting.[8,9] For example, *Bacteroides fragilis* has been linked to colorectal carcinogenesis in its enterotoxigenic form, whereas other *Bacteroides* species have been associated with immune modulation, mucosal homeostasis, or therapeutic responsiveness.[8,10] Recent work has also suggested that *Bacteroides ovatus* (BO) may exert antitumor activity in CRC through metabolite-dependent mechanisms, further raising the possibility that selected commensals may influence tumor behavior in ways not fully captured by conventional immune profiling alone.[10,11]

A major unresolved question is how the microbiome mechanistically alters response to PD-1 blockade. Much of the existing literature has focused on immune-cell priming, antigen presentation, dendritic cell function, and T cell activation.[4,6,12] However, emerging evidence indicates that the microbiome can also shape host and tumor metabolism through microbial products, altered nutrient processing, and systemic metabolic signaling.[12,13] These effects are highly relevant to cancer because metabolic states within the tumor microenvironment influence nutrient availability, redox balance, membrane synthesis, and the functional fitness of both tumor and immune cells, all of which can affect immunotherapy efficacy.[13,14] Recent studies have highlighted microbial metabolites as direct modulators of immunotherapy response, yet whether microbiome-driven metabolic remodeling is a major mechanism linking gut bacteria to PD-1 response in CRC remains insufficiently defined.[12,15] In this study, we sought to define the role of BO in modulating response to PD-1 blockade in CRC and to investigate the underlying metabolic mechanism of BO enhanced immune therapy outcomes. Using a syngeneic CRC tumor model, we demonstrated that BO enhanced the antitumor efficacy of α-mPD-1 therapy. Integrated analyses of fecal microbiota, tumor and serum metabolomics, immune profiling, and multi-omics network interactions revealed that this benefit was independent of major shifts in immune cell composition. Instead, improved therapeutic efficacy was strongly associated with coordinated metabolic remodeling across both local and systemic compartments. Together, our findings support the mechanism whereby BO potentiates immune checkpoint efficacy in CRC by reshaping the microbiome-metabolism axis.

## 2. Materials and Methods

### 2.1 Chemical reagents

*Bacteroides ovatus* (ATCC 8483), used for oral gavage, was obtained from the American Type Culture Collection (ATCC). The MC38 murine colon adenocarcinoma cell line was a gift from Dr. Daniel Spakowicz at The Ohio State University. InVivoMAb anti-mouse PD-1 (α-mPD-1, CD279, Clone: RMP1-14) and its corresponding phosphate-buffered saline (PBS) dilution buffer were purchased from Bio X Cell (Lebanon, NH, USA). Gifu Anaerobic Medium (GAM) for BO culture was purchased from HiMedia (Kennett Square, PA, USA). DMEM, antibiotic–antimycotic solution, 0.25% trypsin–EDTA, and PBS for MC38 cell culture were purchased from Fisher Scientific (Waltham, MA, USA), while fetal bovine serum (FBS) was purchased from Gemini (West Sacramento, CA, USA). QIAamp Fast DNA Stool Mini Kit was purchased from QIAGEN (Germantown, MD, USA) to isolate microbial DNAs from mouse fecal samples for gut microbiome analysis.

### 2.2 Bacterial culture and preparation for oral gavage

BO from a glycerol stock was streaked onto GAM agar plates prepared with 59 g of GAM medium and 15 g of agar per 1 L of deionized water to isolate single colonies for subsequent liquid culture. After inoculation, the plates were incubated at 37 °C for 48 h in a Coy anaerobic chamber. A single colony from the incubated plate was then inoculated into a 50 mL conical tube containing 45 mL of GAM liquid medium (59 g GAM medium per 1 L deionized water) and cultured at 37 °C for 48 h under anaerobic conditions. Optical density (OD) was measured to estimate colony-forming units (CFUs) based on a previously established standard curve generated from drop plate counts. Briefly, a series of dilutions in GAM liquid medium was prepared, and the corresponding OD values were measured to generate a linear calibration curve for CFU estimation in subsequent experiments. Following incubation, BO cells were harvested by centrifugation at 4,000 rpm at 4 °C and resuspended in phosphate-buffered saline (PBS) for oral gavage. Anaerobic conditions in the chamber were maintained using a gas mixture of nitrogen, carbon dioxide, and hydrogen, with oxygen levels maintained between 0 and 10 ppm and hydrogen at 2–3%. Oxygen levels were monitored using a Coy anaerobic monitor (CAM-12). Residual oxygen was removed using two Stak-Pak systems equipped with palladium catalysts, which facilitate the conversion of oxygen to water. A vacuum airlock was used to minimize oxygen exposure during the transfer of reagents and materials into and out of the glove box. In addition, the surfaces of all equipment and materials were disinfected with 10% bleach.

### 2.3 MC38 murine colon adenocarcinoma cell culture

The MC38 murine colon adenocarcinoma cell line was cultured in DMEM supplemented with 10% FBS and 1% antibiotic–antimycotic solution. Cells were maintained at 37 °C in a humidified incubator with 5% CO_2_. Upon reaching approximately 80% confluency, cells were dissociated using 0.25% trypsin– EDTA and 10% of the cells were subcultured. For tumor inoculation, MC38 cells were harvested at ∼80% confluency, washed, and resuspended in phosphate-buffered saline (PBS) at a final concentration of 5 × 10^6^ cells/mL.

### 2.4 Animal studies

#### CRC tumor model construction

For tumor cell implantation, mice were anesthetized with 1–5% isoflurane delivered in oxygen via inhalation and placed in a sterile environment. A suspension of MC38 murine colon adenocarcinoma cells (5 × 10^5^ cells in 100 μL of PBS) was injected subcutaneously into the dorsal flank using a sterile syringe and needle. Following tumor cell injection, animals were monitored until full recovery from anesthesia. Once they were able to stand upright and exhibit normal mobility, they were returned to their home cages. Mice were monitored daily for general health status, discomfort, and clinical distress, and tumor growth was assessed 2–3 times per week by external measurements using calipers. Predefined humane endpoints included severe body weight loss (>15%), tumor ulceration, impaired mobility, or other signs of morbidity. At the experimental endpoint, or earlier if humane endpoints were reached, mice were euthanized by carbon dioxide (CO_2_) inhalation using a University Laboratory Animal Resources (ULAR) euthanasia chamber, followed by removal of vital organs as a secondary method to confirm death. All anesthesia and euthanasia procedures were performed in accordance with the approved Institutional Animal Care and Use Committee (IACUC) protocol (2022A00000054) at The Ohio State University and applicable American Veterinary Medical Association (AVMA) guidelines.

#### Animal housing and experimental design

All animal experiments were conducted in accordance with protocols approved by the Ohio State University Institutional Animal Care and Use Committee (IACUC; Protocol# 2022A00000054). Male C57BL/6 mice (6 weeks old) were obtained from The Jackson Laboratory and housed under controlled conditions (25 °C, 12-h light/dark cycle). A total of 60 mice were used across two experimental cohorts: metabolomics/microbiome (n = 40) and flow cytometry (n = 20). Following a one-week acclimation period, mice were assigned to four treatment groups (n = 10 or 5 per group, depending on the cohort). Baseline blood and fecal samples were collected prior to cell injection. Mice were then subcutaneously injected with MC38 cells. On day 3 post-inoculation, mice received oral gavage of 5 × 10^7^ BO cells, with PBS administered to control groups. BO administration was repeated weekly. When tumors became palpable (approximately day 8 post-inoculation), mice were treated with 100 µg α-mPD-1 per mouse via intraperitoneal injection once weekly, with PBS used as a control. Tumor growth was monitored by digital caliper measurements at designated time points. Individual mouse tumor-volume (= 0.5 x length x width^2^) measurements used for longitudinal and endpoint analyses are provided in **Table S1-1**. At study endpoints (day 16), mice were euthanized. Tumors were aseptically excised following surface sterilization with 70% ethanol. Blood samples were collected and centrifuged at 1,500 rpm to isolate serum for metabolomics analysis. Fecal samples were also collected immediately prior to euthanasia.

### 2.5 16S rRNA gene sequencing analysis

Bacterial community composition in the fecal samples was characterized by sequencing the V4 region of the 16S rRNA gene from fecal DNA as we have described in our previous publication.[16] Amplicon libraries were prepared using the 515F/806R primer pair and barcoded primers incorporating adapter sequences compatible with Illumina sequencing. PCR amplification was performed in a total reaction volume of 25 µL consisting of 9.5 µL of certified DNA-free PCR water (MO BIO), 12.5 µL of AccuStart II PCR ToughMix (QuantaBio; 2× stock, 1× final concentration), 1 µL each of the Golay barcode-tagged forward primer and reverse primer (5 µM stocks; 200 nM final concentration), and 1 µL of template DNA. Thermal cycling began with an initial denaturation at 94 °C for 3 min, followed by 35 cycles of 94 °C for 45 s, 50 °C for 60 s, and 72 °C for 90 s. A final extension was performed at 72 °C for 10 min. Following amplification, PCR products were quantified using the PicoGreen assay (Invitrogen) on an Infinite 200 PRO plate reader (Tecan). Amplicons from individual samples were subsequently combined in equimolar proportions to generate a single pooled library. The pooled library was purified with AMPure XP beads (Beckman Coulter) and its concentration was measured using a Qubit fluorometer (Invitrogen). Based on the resulting concentration, the library was adjusted to 2 nM, denatured, and diluted to a final loading concentration of 6.75 pM. PhiX control library was added at 10% before sequencing. Sequencing was conducted on an Illumina MiSeq using a 151 bp × 12 bp × 151 bp run configuration with customized sequencing primers and procedures. The resulting 16S rRNA gene amplicon sequencing data were processed and analyzed using QIIME 2 through Ohio Supercomputer Center (OSC).

### 2.6 Culturomic expansion and qPCR detection of intratumoral *Bacteroides ovatus*

To expand translocated intratumoral bacteria from the PBS, α-mPD-1, BO, and BO + α-mPD-1 groups, we performed culturomics as described previously.[17] Briefly, tumor tissues were aseptically collected and homogenized in 0.05% NP-40 (IGEPAL; Sigma-Aldrich, I8896) by mashing the tissues through 40-μm filters into 50-mL conical tubes. The resulting tumor homogenates were incubated overnight at 37 °C under anaerobic conditions. Following incubation, the tumor homogenate cultures were collected, and genomic DNA was extracted using the Wizard Genomic DNA Purification Kit (Promega, Madison, WI, USA). To assess the presence of BO, quantitative PCR (qPCR) was performed using the extracted DNA with BO-specific primers (forward, 5′-TGCAAACTRAAGATGGC-3′; reverse, 5′-CAAACTAATGCAACGCATC-3′) and universal 16S rRNA gene primers (forward, 5′-ACTCCTACGGGAGGCAGCAG-3′; reverse, 5′-ATTACCGCGGCTGCTGG-3′). For each technical replicate, ΔCt was calculated as Cq(BO target)−Cq(16S). ΔΔCt was subsequently calculated as ΔCt(sample)−mean ΔCt(PBS), and the relative BO signal was determined using the 2^−ΔΔCt^ method. The resulting qPCR data are provided in **Table S2** and were used for subsequent statistical analyses.

### 2.7 Metabolites extraction from tumor, mice serum and fecal pellets

Metabolite extraction procedures were modified based on we previously described.[18,19] Briefly, tumor and fecal pellets samples were placed in 2 mL screw-cap tubes containing three 2.7 mm glass beads, while 20 μL of mouse serum was added directly to 1.5 mL centrifuge tubes without glass beads. Methanol (250 μL) and 50 μL of a ^13^C^15^N-labeled amino acid internal standard mixture were added to all samples. Tumor samples were homogenized using a BioSpec Mini-BeadBeater for three 15-second cycles. Both homogenates were incubated at −20 °C for 30 min to precipitate proteins and insoluble material, followed by centrifugation at 14,000 rpm for 20 min at 4 °C. Supernatants (150 μL) were transferred to fresh tubes and dried in an Eppendorf SpeedVac at 30 °C for approximately 2 h. Dried extracts were reconstituted in 250 μL of 50% acetonitrile, vortexed for 2 min, and centrifuged at 14,000 rpm for 5 min at 4 °C. A 200 μL aliquot of the clear supernatant was used for high-Performance liquid chromatography–high-resolution mass spectrometry (HPLC–HRMS) analysis. For quality control (QC), 10 μL of supernatant from each sample was pooled for compound annotation using data-dependent acquisition (DDA) mode (top 10 precursor ions).

### 2.8 Detection of metabolites in ultra-high performance liquid chromatography coupled to high-resolution mass spectrometry (UHPLC-HRMS)

Polar metabolite profiling of tumor, serum, and fecal extracts was carried out using hydrophilic interaction liquid chromatography coupled with high-resolution mass spectrometry. Samples were injected onto an XBridge BEH Amide column (2.5 µm, 2.1 × 150 mm; Waters, Milford, MA, USA) operated at 40 °C on a Thermo Vanquish UPLC system. The polar mobile phase (A) contained water/acetonitrile (90:10, v/v), whereas the nonpolar mobile phase (B) contained water/acetonitrile (10:90, v/v); both phases were supplemented with 5 mM ammonium acetate and 0.1% acetic acid. Chromatographic elution was performed at 0.3 mL/min over 12 min. The proportion of solvent B was initially set at 70%, reduced to 30% over the first 5 min, maintained at 30% through 8 min, increased back to 70% by 9.5 min, and held at this composition for the remaining of the run to re-equilibrate the column.[20] Eluting metabolites were detected with a Q Exactive Hybrid Quadrupole-Orbitrap mass spectrometer (Thermo Fisher Scientific, Waltham, MA, USA) operated separately under positive- and negative-ion conditions. Full MS-SIM acquisition was used with an Orbitrap resolution of 70,000, an AGC target of 3 × 10^6^, a maximum ion injection time of 200 ms, a chromatographic peak-width setting of 6 s, and a mass range of m/z 60–900. For positive-ion acquisition, the electrospray source was configured with sheath, auxiliary, and sweep gas settings of 50, 15, and 1, respectively; the spray voltage was 3.5 kV, the capillary temperature was 350 °C, the S-lens RF level was 50, and the auxiliary gas heater temperature was 425 °C. The same settings were applied for negative-ion acquisition, with the exception of the spray voltage, which was reduced to 2.75 kV.[20]

### 2.9 Tumor infiltrating leukocytes (TIL) isolation and flow cytometry

Tumor tissues were digested in RPMI with 2% FBS (Gemini Bio) containing DNase I (1 μg/mL, Sigma-Aldrich) and collagenase IV (1 mg/mL, Sigma-Aldrich), followed by digestion at 37 °C for 1 h and filtration through a 70-μm cell strainer. Cell suspensions were subjected to ACK lysis and Percoll density gradient separation (800g, 30min, 25 °C) using 40% and 80% Percoll to enrich TILs. For surface staining, cells were resuspended in fluorescence-activated cell sorting (FACS) buffer (1% BSA, 1 mM EDTA, 0.05% sodium azide) and blocked with anti-CD16-32 antibody (clone 2.4G2; 5 µg/mL; BioXCell) for 10 min on ice. The fluorescence conjugated Abs and live-dead dye were added and incubated for another 25-30 min. Cells were washed two times with FACS buffer, fixed with 1% paraformaldehyde (Thermo Fisher) and analyzed by BD symphony or Cytek Aurora. For intracellular cytokine staining, 1 × 10^6^ cells were stimulated in complete RPMI containing phorbol 12-myristate 13-acetate (10 nM; Sigma-Aldrich) ionomycin 500 nM; Sigma-Aldrich) and Brefeldin A (1,000×; Biolegend) at 37 °C with 5% CO_2_ for 3.5–4 h. After surface staining, cells were fixed with 1× True Nuclear fixation Buffer (Biolegend) and washed with 1× True Nuclear permeabilization buffer. Cells were stained with fluorescence conjugated Abs on ice for 30 min. After additional washes, cells were resuspended in FACS buffer and analyzed by flow cytometry.

### 2.10 Data processing

#### Metabolites identification

Spectra of polar metabolites were processed using Compound Discoverer 3.3 (Thermo Fisher Scientific, Waltham, MA, USA) for compound identification. The resulting dataset was exported as an Excel file containing metabolite name, database source, m/z, retention time (RT), and peak area for each detected feature. Unidentified features were excluded. The remaining metabolites were matched against in-house reference libraries mzCloud and mzVault, and in-house data library MassList, which had been validated by collecting spectra on the same instrument with commercial analytical chemical standards. Annotated compounds were further filtered to retain only those with pooled QC covariance (CV) < 25%. After duplicate removal, the final metabolite set (**Table S3-1 for tumor samples, Table S4-1 for serum samples, and Table S5-1 for fecal samples**) was used for downstream analyses in MetaboAnalyst 6.0 and R 4.3.0.[20]

#### 16S rRNA data processing

Paired-end 16S rRNA gene amplicon reads were processed in QIIME 2 (v2023.7). Raw FASTQ files were imported using a Phred33-formatted paired-end manifest, and sequence quality was assessed before denoising. Amplicon sequence variants (ASVs) were inferred using DADA2 with truncation lengths of 220 bp for forward reads and 155 bp for reverse reads, with pooled chimera removal enabled. Representative ASV sequences were taxonomically assigned using a pre-trained SILVA 138 99% classifier targeting the 515F/806R region. Features annotated as mitochondria, unassigned taxa, or uncultured organisms were removed before downstream analysis, and taxonomic composition was visualized with taxonomic bar plots. The final data table used for downstream analysis was with ASV and taxonomic information was reported in **Table S6-1** and **Table S6-2.**

### 2.11 Data analysis and visualization

All statistical tests were two-sided. The effects of BO and α-mPD-1 treatment were evaluated using a 2 × 2 factorial designs, with BO and α-mPD-1 included as binary factors. Linear regression models were used to estimate the main effects of each treatment and their interaction (BO × α-mPD-1). The interaction term was used to assess whether the combined treatment effect differed from the additive effects of the individual treatments on the model’s analysis scale as the following:

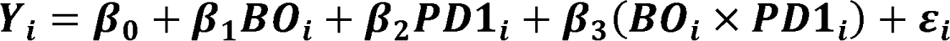

Where *Y_i_* represents the tumor burden outcome for mouse *i*, *Bo_i_* and *PD1_i_* are centered treatment indicators coded as −0.5 (untreated) and +0.5 (treated), respectively. ε*_i_* denotes the residual error. The intercept β*_0_* represents the equally weighted mean outcome across the four treatment groups, while β*_1_* and β*_2_* estimate the factorial main effects of BO and α-mPD-1, respectively. The interaction coefficient β*_3_* evaluates whether the effect of α-mPD-1 differs according to BO treatment status on the analysis scale. Heteroscedasticity-consistent (HC) standard errors were estimated using the HC3 covariance estimator, and all statistical tests were two-sided. Tumor-growth analyses began with observed measurements on day 8; visualization-only values at days 0, 3, and 6 were excluded from inference. Tumor volume *Y_i_* was analyzed as log (1 + tumor volume). Time-averaged burden over days 8–14 was calculated from trapezoidal AUC/6 and analyzed together with day-14 and available-case day-16 endpoints using HC3 models. Longitudinal tumor growth was analyzed using Gaussian generalized estimating equations (GEE), with day treated as a categorical variable and an exchangeable working correlation structure assuming a common correlation between repeated measurements within each mouse. Day-16 missing values were not imputed. Microbiome analyses were performed at the genus level. Relative abundance was calculated by closure, Shannon diversity used natural logarithms, and Centered Log Ratio (CLR) transformation used a 0.5-read pseudocount. Paired baseline-to-final changes were analyzed within mouse. Bray–Curtis and Aitchison distances were evaluated by Principal Coordinates Analysis (PCoA), Permutational Multivariate Analysis of Variance (PERMANOVA), and Permutational Analysis of Multivariate Dispersions (PERMDISP) using 9,999 permutations, with sensitivity analyses after removal of Bacteroides. Genus-level differential abundance used paired CLR changes, HC3 models, and Benjamini–Hochberg (BH) correction. Tumor metabolomics was evaluated in 16 mice, four per treatment group, after averaging three technical UHPLC-HRMS injections per mouse on the normalized log_2_ scale. Technical-precision QC retained 226 of 243 detected features (**Table S3-2**). Feature-wise effects were analyzed by 2 × 2 HC3 models with BH correction. Four prespecified tumor metabolic modules: glycan turnover, activated donors, oxidized lipids, and fatty-acid trafficking, were summarized as mean z scores of their constituent metabolites. Serum and fecal metabolomics were analyzed as within-mouse final-minus-baseline changes; serum included 39 paired mice and 167 features, whereas fecal metabolomics included 19 paired mice and 303 features and was considered exploratory. Associations among microbial abundance, metabolic features, metabolic modules, and tumor burden were evaluated using treatment-adjusted partial Spearman correlations. To account for treatment-related differences, each pair of variables was independently rank-transformed and regressed against BO treatment, α-mPD-1 treatment, and their interaction. Pearson correlations between the resulting residuals were then calculated to estimate associations independent of the modeled treatment-group effects. For feature-wide associations involving *Bacteroides* and for metabolic module– tumor burden associations, P values were adjusted for multiple testing using the BH method within predefined testing groups. An integrated correlation network was constructed using 25 preselected features representing microbial, metabolic, and tumor-related variables. Correlation testing was restricted to pairs of features from different data categories, with at least 16 mice having valid measurements for both features. Network edges were displayed when the unadjusted P value was < 0.05 to visualize exploratory associations. All analyses and figures were generated using R 4.6.0.

## 3. Results

### 3.1 Successful colonization of *Bacteroides* reshapes the gut microbial community

To determine whether BO successfully colonized the gut and altered fecal microbial composition during treatment, 16S rRNA gene sequencing was performed on fecal samples collected at baseline and at the experimental endpoint (**Figure 1**). The genus-level abundance and taxonomic data underlying these analyses are provided in **Table S6-1** and **Table S6-2**. Genus-level microbiome composition shifted visibly in both BO-administered groups, with *Bacteroides* comprising on average 34.62% of final abundance of BO and 34.51% in BO + α-mPD-1, compared with 0.075% in PBS and 0.120% in α-mPD-1 (**Figure 1A-1B**). CLR of *Bacteroides* increased within every group, but the changes were far larger after BO administration: with a mean change of 9.520 CLR units in BO and 9.558 in BO + α-mPD-1, while the paired-change interaction was not significant (q=0.991) (**Figure 1B and Table S6-3**). Shannon diversity decreased significantly from baseline to final sampling in the PBS and BO groups, while smaller and nonsignificant decreases were observed in the α-mPD-1 (mean ΔShannon = −0.131, P = 0.384) and BO + α-mPD-1 (mean ΔShannon = −0.121, P = 0.384) groups (**Figure 1C and Table S6-3**). At the microbiome community level, final Bray–Curtis profiles differed among groups, while the corresponding dispersion test was not significant (p=0.310) (**Figure 1D and Table S6-3**). Within-mouse baseline-to-final Bray–Curtis distance also differed by group; BO + α-mPD-1 exceeded α-mPD-1 by 0.143 distance units (**Figure 1E and Table S6-3**). Removing *Bacteroides* did not abolish treatment-associated community structure. Bray–Curtis PERMANOVA without *Bacteroides* remained significant, with no dispersion difference (P=0.310) (**Figure S1 and Table S6-3**). In paired-CLR differential abundance, 7 of 35 genera met BH q<0.05 for the average BO effect (**Figure S2 and Table S6**). *Bacteroides* and the displayed A2 and *Monoglobus* features increased, whereas *Dubosiella*, *Tyzzerella*, *Lachnospiraceae* NK4A136 group, and *Acetatifactor* decreased (**Figure S2 and Table S6-4**). These changes indicate that BO exposure coincided with broader genus-level microbiome remodeling rather than only expansion of the administered *Bacteroides*.

**Figure 1.**
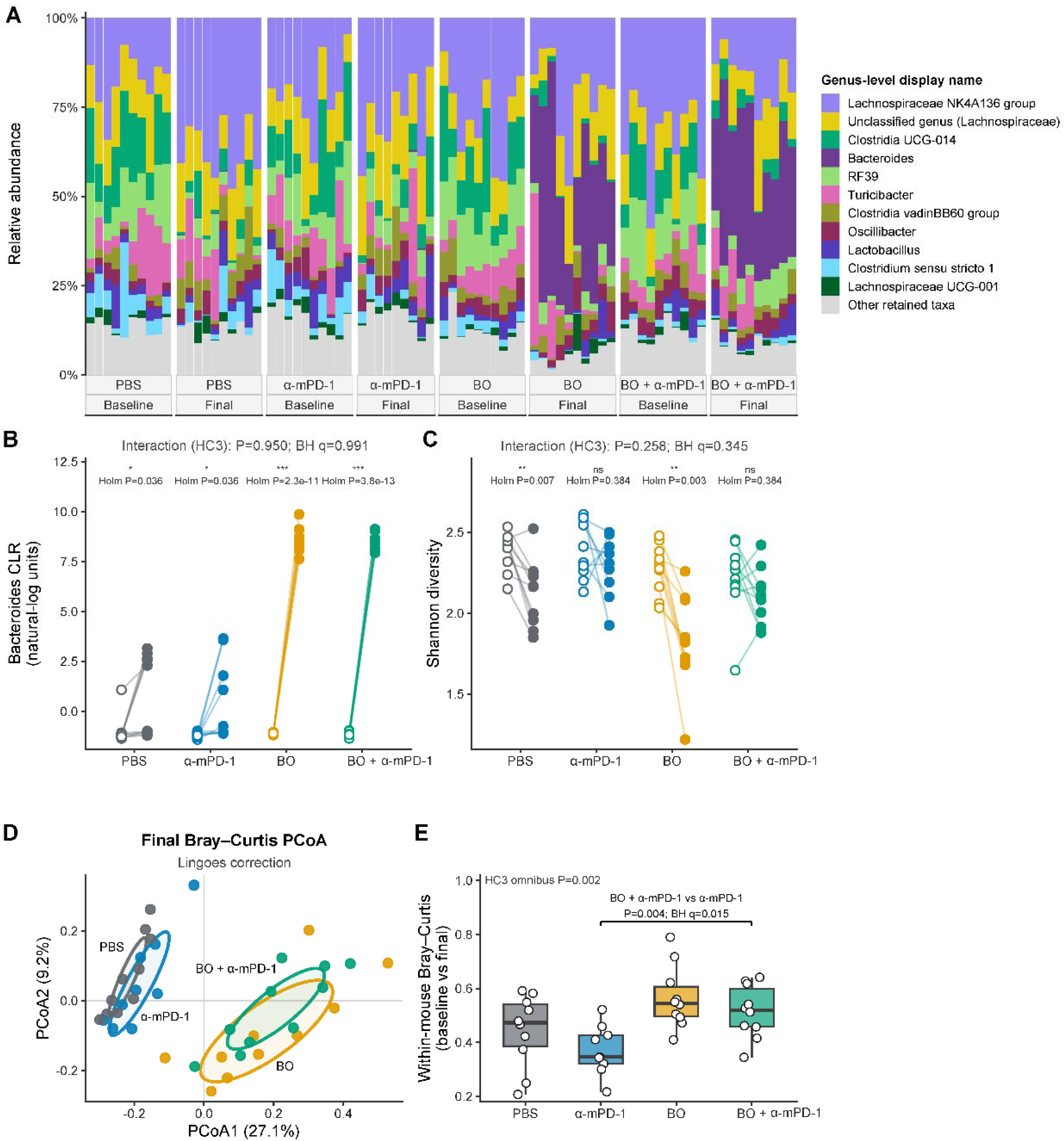
Longitudinal genus-level microbiome remodeling. Paired baseline and final fecal 16S profiles from 39 mice: PBS n=10, α-mPD-1 n=9, BO n=10, and BO + α-mPD-1 n=10. **(A)** Individual-sample relative-abundance bars after aggregation by the explicit genus field. The 11 most abundant genus-level display names are colored and remaining retained genera are pooled as ‘Other retained taxa.’ Unresolved genera use the audited ‘Unclassified genus (nearest resolved parent)’ convention. **(B)** Paired *Bacteroides* CLR values, calculated after adding a 0.5-read pseudocount. **(C)** Paired Shannon diversity. In both Panels B and C, open and filled points denote baseline and final samples, respectively, and lines connect mice. The interaction statistic shown in Panels B and C belongs to a separate factorial hypothesis family and is displayed as a visually secondary annotation. **(D)** Final-sample Bray–Curtis PCoA with Lingoes correction. Ellipses are 95% Hotelling T^2^ confidence regions for treatment-group centroids, calculated from the displayed two-dimensional PCoA coordinates under a bivariate-normal approximation; they are visualization aids only and were not used in PERMANOVA or PERMDISP. The locked final-sample tests were PERMANOVA R^2^=0.269; raw P=0.0001; BH q=0.00015; PERMDISP raw P=0.212; BH q=0.310. **(E)** Within-mouse Bray–Curtis distance between baseline and final profiles. Boxes show median and IQR, whiskers 1.5×IQR, and points are mice.

### 3.2 Administration of BO slowed tumor growth

Tumor growth was evaluated in 38 mice with complete observations from day 8 through day 16 (**Table S1-1**). Zero values shown on days 0, 3, and 6 represent the pre-palpable phase and were included for visualization purposes only and were excluded from all statistical analyses. Given the heterogeneous response to α-mPD-1 monotherapy, with both responders and nonresponders observed in this group, the subsequent analyses primarily focused on comparisons between the BO + α-mPD-1 combination group and the PBS control group. Mean ± SEM growth trajectories indicated that tumors in the BO+α-mPD-1 group exhibited delayed tumor progression relative to the PBS group after treatment initiation and remained the smallest among all treatment groups through the end of the complete-data window (**Figure 2A**). For time-averaged tumor burden across days 8-14, the BO+α-mPD-1 group exhibited a lower tumor burden than the PBS group, with an estimated effect of −0.293 log1p(mm³), corresponding to a 25.4% reduction in the geometric mean of 1+tumor volume (ratio = 0.746, **Figure 2B** and **Table S1-2**). By day 14, the treatment effect had increased to −0.751 log1p(mm³), corresponding to a geometric mean ratio of 0.472, or an approximately 52.8% lower tumor burden relative to PBS (**Figure 2C and Table S1-2**). Although the BO+α-mPD-1 group exhibited the lowest tumor burden, factorial analyses did not provide evidence for synergy. BO × α-mPD-1 interaction terms were not significant for either time-averaged tumor burden (days 8-14; q = 0.724) or day-14 tumor burden (q = 0.724) (**Table S1-2**). Repeated-measures GEE analyses similarly detected no significant day × BO × α-mPD-1 interaction (q = 0.418) (**Table S1-2**). Descriptive analysis of available day-16 data showed the lowest mean tumor volume in the BO+α-mPD-1 group (76.53 mm³), followed by BO (122.17 mm³), α-mPD-1 (145.94 mm³), and PBS (189.12 mm³), consistent with additive antitumor effects of the combined treatment (**Figure 2D and Table S1-2**). Available-case analyses at day 16 further supported the antitumor benefit of BO+α-mPD-1, which remained significantly lower than both PBS controls and α-mPD-1 alone. Nevertheless, the BO × α-mPD-1 interaction was not significant, indicating that the enhanced tumor suppression observed with combination treatment did not meet statistical criteria for synergy (**Table S1**). Overall, BO+α-mPD-1 achieved the greatest reduction in tumor burden throughout the study, supporting enhanced therapeutic efficacy of the BO+α-mPD-1 treatment while indicating predominantly additive, rather than synergistic, antitumor effects.

**Figure 2.**
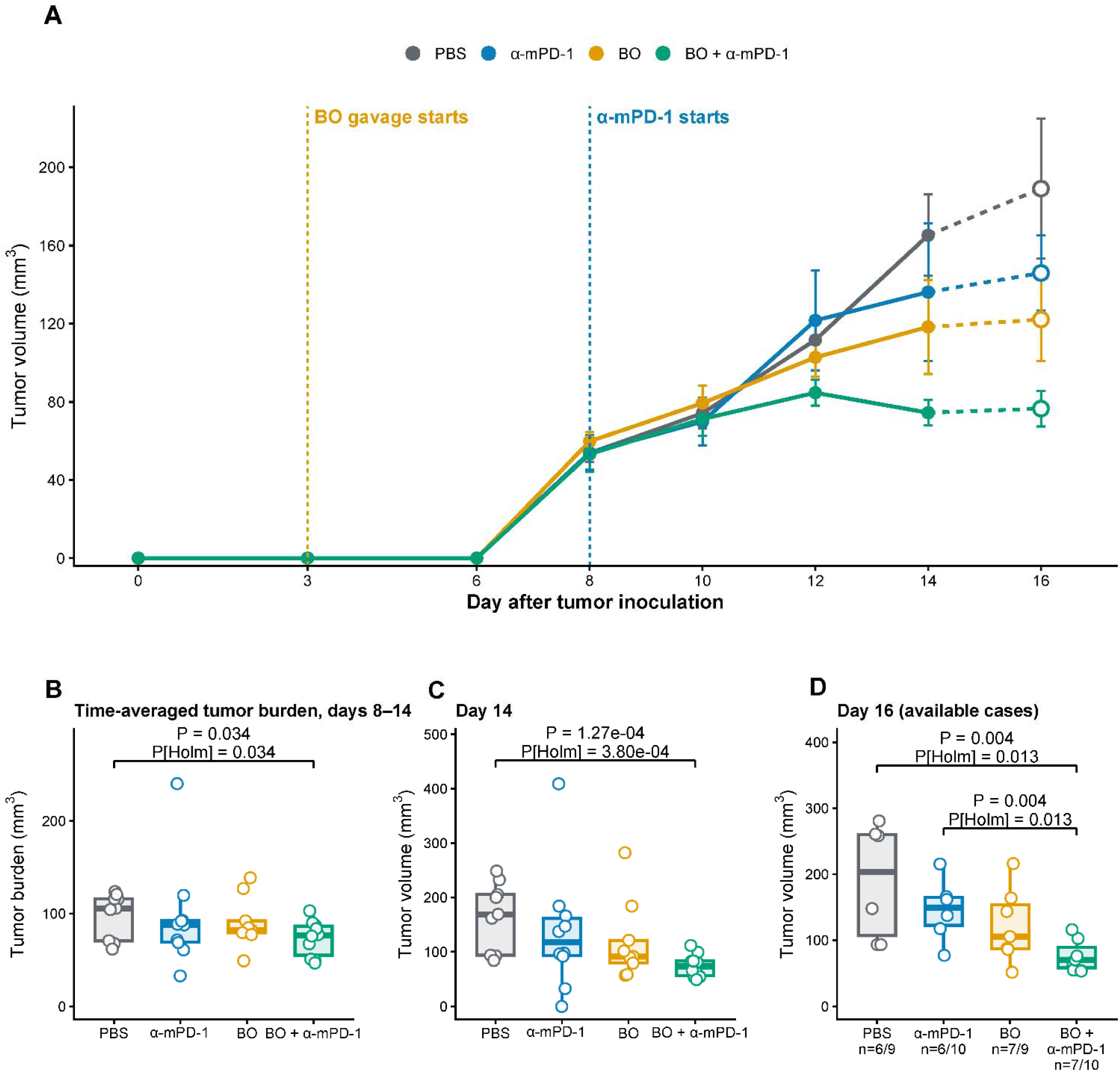
BO+α-mPD-1 treatment produced the strongest observed tumor-control phenotype. **(A)** Group mean ± SEM tumor-volume trajectories for PBS (n=9), α-mPD-1 (n=10), BO (n=9), and BO + α-mPD-1 (n=10). MC38 cells were implanted on day 1, BO gavage began on day 3, and α-mPD-1 began on day 8. Day 1, 3, and 6 values are visualization-only zeros; inference begins with observed day-8 values. The day 14–16 connectors are dashed and day-16 markers are open because only 26/38 day-16 measurements were available; no imputation was used. **(B)** Mouse-level time-averaged tumor burden, days 8–14. Time-averaged burden was calculated as trapezoidal AUC from Days 8–14 divided by the six-day interval. **(C)** Mouse-level day-14 tumor volume. **(D)** Day-16 available cases, with observed/total n printed below each group. Boxes show median and interquartile range, whiskers extend to 1.5×IQR, and points are mice. Endpoint models used log1p(mm³), a 2 × 2 factorial design, and HC3 covariance.

### 3.3 BO was enriched in tumor-derived cultures following BO administration

To assess whether BO could be detected in tumor-derived bacterial cultures, DNA extracted following anaerobic expansion of tumor homogenates was analyzed by BO-specific qPCR and normalized to the universal 16S rRNA gene signal. BO-treated groups exhibited substantially lower ΔCt values than the PBS and α-mPD-1 groups, indicating a markedly greater abundance of BO-associated sequences relative to total bacterial 16S signal (**Figure S3A; Table S2**). Consistent with this pattern, normalization to the PBS group using the 2^−ΔΔCt^ method revealed a strong enrichment of the relative BO-associated signal in both the BO and BO+α-mPD-1 groups, reaching approximately 10^4^-fold above the PBS reference, whereas the α-mPD-1 group showed little BO-associated signal (**Figure S3B; Table S2**). Together, these findings demonstrate marked enrichment of BO-associated DNA in tumor-derived bacterial cultures from BO-treated mice, supporting the presence of BO in tumors following oral administration.

### 3.4 BO+α-mPD-1 treatment induces broad tumor-metabolic remodeling

Metabolic reprogramming in CRC has been observed from previous studies, and dysregulated fatty acid (FA) metabolism is an established driver of both tumor cell proliferation and immune evasion.[21–24] Therefore, we next performed global tumor metabolomic profiling using UHPLC-HRMS to identify treatment-induced metabolic changes in mice tumor tissues. In total, we detected 243 metabolites (**Table S3-1)**. PCA of the QC-passing mouse-level matrix showed treatment-induced changes in metabolic profiles despite limited dispersion difference (q=0.393) (**Figure 3A; Figure S4)**. Compared with PBS, BO + α-mPD-1 treatment was associated with significant differences in 135 of 226 metabolic features at the nominal P < 0.05 threshold, of which 130 remained significant after BH correction (q < 0.05; **Figure 3B; Table S3-3**). In contrast, factorial interaction analysis identified 32 features with nominally significant BO × α-mPD-1 interactions (P < 0.05), but only hydroxylysine remained significant after multiple-testing correction (q < 0.05; **Figure S4D**). To better capture coherent biochemical patterns, we prespecified four modules based on the significant changed metabolites from above analysis: glycan-turnover related metabolites, activated donors, oxidized lipids, and fatty-acid trafficking metabolites (**Figure 3C–G**). Relative to PBS, the BO+α-mPD-1 group had a higher glycan-turnover score, lower activated-donor score, higher oxidized-lipid score, and higher fatty-acid-trafficking score (**Figure 3D– G; Table S3-4**). Formal interaction q values were 0.402, 0.402, 0.402, and 0.221, respectively (**Figure 3D–G; Table S3-4)**. Thus, the BO+α-mPD-1 group showed strong module-level differences from PBS, but the modules did not meet the statistical criterion for BO × α-mPD-1 synergy. The glycan-turnover module comprised iditol, mannose, sucrose, fucose, and stachyose, and the fucose and iditol were significantly increased in the BO+α-mPD-1 versus PBS comparison (**Figure 3C**). Citicoline and CDP-ethanolamine, the two activated-donor metabolites, were both significantly lower in the BO+α-mPD-1group versus PBS (**Figure 3C**). The oxidized-lipids module contained 15-HEPE and 12-HPETE; 12-HPETE was observed to significantly higher in BO+α-mPD-1 compared to PBS group (**Figure 3C**). The fatty-acid-trafficking module comprised three acylcarnitines (palmitoylcarnitine, propionylcarnitine, and acetylcarnitine) and two dicarboxylic acids (sebacic acid and dodecanedioic acid). Among these, palmitoylcarnitine, sebacic acid, and dodecanedioic acid showed significantly higher abundances in the BO + α-mPD-1 group than in PBS controls after FDR correction (BH-adjusted q < 0.05; **Figure 3C and Table S3-3**).

**Figure 3.**
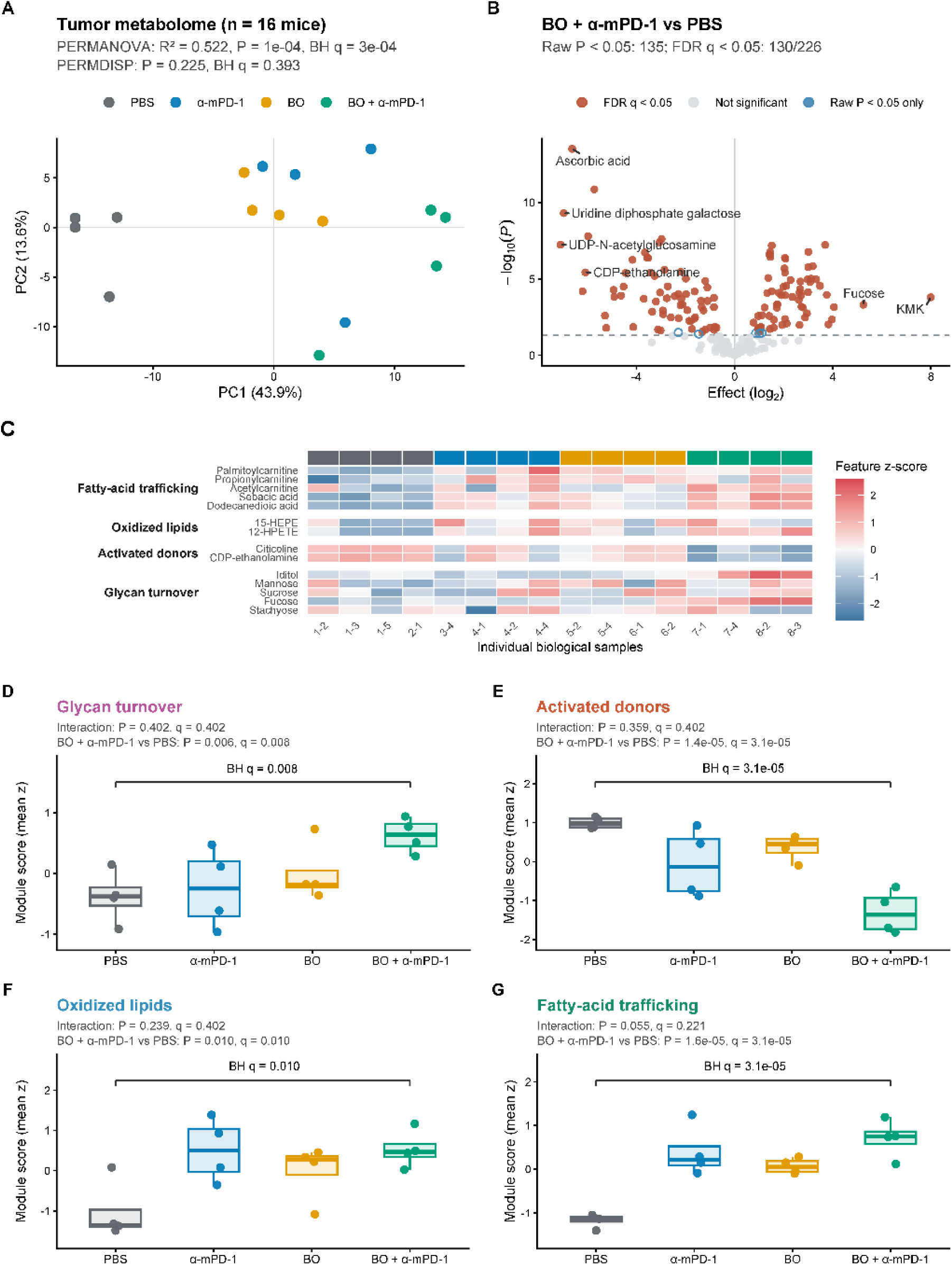
Tumor-metabolomic remodeling in the 2 × 2 treatment design. Tumor metabolomics used 16 mice (n=4 per group). Each tumor had three technical injections; technical injections were median-normalized and log_2_ transformed, then averaged to one mouse-level value. Features passed QC when median technical CV≤25% and ≤25% of mice had CV>30%; 226/243 features were retained. **(A)** PCA of the 226-feature mouse-level matrix. The locked global tests were PERMANOVA R^2^=0.5216; raw P=0.0001; BH q=0.0003; PERMDISP raw P=0.225; BH q=0.393. **(B)** Volcano plot for BO + α-mPD-1 versus PBS from HC3 factorial contrasts. Filled red points have BH q<0.05, open blue points have raw P<0.05 only, and gray points do not meet raw P<0.05; the horizontal line marks P=0.05. **(C)** Heatmap of 14 detected prespecified module features; columns are biological mice and cells are feature z scores. Technical replicates are not displayed as independent samples. **(D–G)** Mouse-level module scores for glycan turnover, activated donors, oxidized lipids, and fatty-acid trafficking. Scores are means of member-feature z scores. Boxes show median and IQR, whiskers 1.5×IQR, and points are mice.

To investigate whether the improved antitumor response associated with BO administration was accompanied by changes in the tumor immune microenvironment, we performed flow cytometric analysis of tumor-infiltrating immune cells on day 14. Overall, no consistent treatment-associated differences were detected in the major immune cell populations examined (**Figure S5**). The frequencies of CD4^+^ and CD8^+^ T cells were broadly comparable across treatment groups, although CD8^+^ T-cell frequency was numerically lower in the BO + α-mPD-1 group. Similarly, no consistent differences were observed in activation-associated (CD44^hi^), exhaustion-associated (PD-1^+^Tim-3^+^), or memory-associated (TCM, TEM, and TN) T-cell phenotypes within the CD4^+^ and CD8^+^ compartments (**Table S7; Figure S5B-D**). Macrophage frequency was modestly higher in the BO + α-mPD-1 group than in PBS controls but was comparable to the other treatment groups (**Figure S5A**). No consistent differences in M1-like or M2-like macrophage polarization were observed (**Figure S5C**). Together, these findings suggest that the improved antitumor response associated with BO + α-mPD-1 treatment was not accompanied by substantial or consistent changes in the measured immune cell populations at day 14. In contrast, tumor metabolomic profiling revealed pronounced treatment-associated metabolic alterations, including changes in glycan turnover, activated-donor metabolism, oxidized lipids, and fatty-acid-associated metabolites. These findings suggest that metabolic remodeling may contribute to the antitumor response associated with BO administration and PD-1 blockade, although further functional studies are required to establish the underlying mechanisms.[25–27]

### 3.5 Serum and fecal metabolomics identified nominal circulation and gut level metabolic changes induced by BO+α-mPD-1 treatment

We next performed serum and fecal metabolomics analyses to explore systematic metabolic features that are related to the treatment. Serum metabolomics analyses detected 167 annotated metabolic features, whereas fecal dataset detected 303 metabolic features; complete feature-level results are provided in **Tables S4-1 and S5-1**, respectively. PCA showed treatment-associated multivariate structure, whereas dispersion did not differ detectably (PERMDISP P=0.262; BH q=0.393) (**Figure 4A; Table S5-3**). In the BO + α-mPD-1 versus PBS comparison, 25 of 167 serum metabolic features showed nominally significant differences in baseline-to-final changes (P < 0.05), but only xanthosine remained significant after BH correction (**Figure 4B; Table S4-2**). Xanthine, deoxyribose 5-phosphate, tyramine, hippuric acid, and cholic acid also exhibited nominally significant differences (P < 0.05), but none survived FDR correction. These findings suggest treatment-associated changes in systemic metabolic profiles, with xanthosine showing the strongest statistical evidence in the prespecified comparison. Due to limited fecal sample availability, fecal metabolomic analyses were considered exploratory. No significant differences in overall metabolic profiles were detected among treatment groups (PERMANOVA, R² = 0.184, P = 0.290), and group dispersion did not differ significantly (PERMDISP, P = 0.751; **Figure 4C; Table S5-3**). In the BO + α-mPD-1 versus PBS comparison, 44 of 303 metabolic features showed nominally significant differences (P < 0.05), but none remained significant after BH correction (**Figure 4D; Table S5-2**). In the α-mPD-1 versus PBS comparison, acetylcholine showed a significantly lower baseline-to-final change but not in other comparisons (**Table S5-2**). Given the small sample size, these findings were considered exploratory. Raw total metabolite signal changes between baseline and final samples are shown in **Figure S7**.

**Figure 4.**
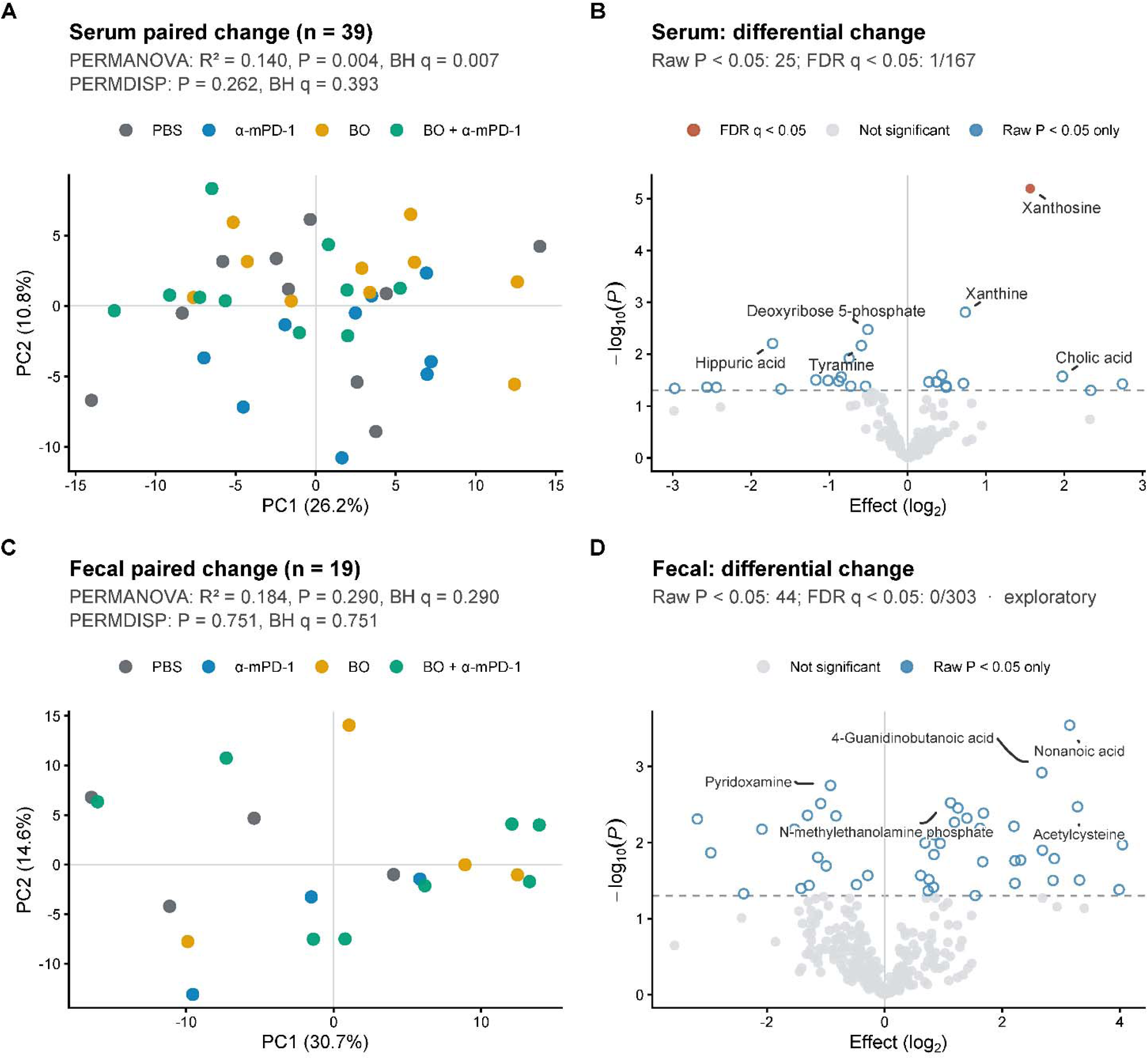
Longitudinal serum and fecal metabolomic changes. Values are within-mouse final-minus-baseline changes in log median-normalized intensity. **(A)** PCA of serum paired changes (n=39; PBS 10, α-mPD-1 9, BO 10, BO + α-mPD-1 10) across 167 features. **(B)** Serum BO + α-mPD-1 versus PBS volcano plot from HC3 factorial models. **(C)** PCA of fecal paired changes (n=19; PBS 4, α-mPD-1 3, BO 4, BO + α-mPD-1 8) across 303 features. **(D)** Fecal BO + α-mPD-1 versus PBS volcano plot; this analysis is exploratory because of small unequal groups. The locked serum global tests were PERMANOVA R^2^=0.140; raw P=0.0044; BH q=0.0066; PERMDISP raw P=0.262; BH q=0.393; the locked fecal global tests were PERMANOVA R^2^=0.184; raw P=0.290; BH q=0.290; PERMDISP raw P=0.751; BH q=0.751. Volcano colors distinguish BH q<0.05, raw P<0.05 only, and nonsignificant features; horizontal lines mark P=0.05.

### 3.6 Integrated analysis reveals coordinated tumor metabolic remodeling beyond treatment-group effects

We next investigated associations among *Bacteroides* abundance, tumor metabolic modules, and day-14 tumor volume before and after adjustment for treatment-group effects (**Figure 5A and Figure 5B**). In unadjusted analyses, the activated-donor module showed a strong inverse correlation with oxidized lipids (ρ = −0.809, q = 0.0011) and fatty-acid trafficking (ρ = −0.921, q < 0.001), while showing a positive correlation with day-14 tumor volume (ρ = 0.621, q = 0.035). Oxidized lipids and fatty-acid trafficking were also positively correlated (ρ = 0.762, q = 0.0030; **Figure 5A; Table S8**). *Bacteroides* abundance showed nominal correlations with glycan turnover and day-14 tumor volume, but neither survived FDR correction. After adjusting for BO, α-mPD-1, and their interaction, the negative correlations between the activated-donor module and both oxidized lipids (partial ρ = −0.741, q = 0.028) and fatty-acid trafficking (partial ρ = −0.851, q = 0.0034) remained significant (**Figure 5B and Table S8**). These findings indicate that associations among these metabolic modules persisted after accounting for treatment-group differences. In contrast, correlations between metabolic modules and day-14 tumor volume were attenuated and were no longer significant after adjustment (all q ≥ 0.742). Consistent results were obtained in the dedicated module-tumor correlation analyses (**Table S8**). Before treatment adjustment, glycan turnover, activated donors, oxidized lipids, and fatty-acid trafficking showed varying degrees of association with tumor volume (**Figure 5A and Table S8**). These module–tumor correlations were substantially attenuated after adjustment for treatment-group effects, suggesting that the unadjusted associations were partly attributable to differences among treatment groups rather than independent variation within groups. A similar pattern was observed in feature-wide analyses of *Bacteroides* abundance and individual metabolites. Among 226 tumor metabolic features and 167 serum paired-change features, none showed significant treatment-adjusted correlations with *Bacteroides* after BH correction (all q ≥ 0.10; **Figure 5C-D; Table S8**). Among 303 fecal paired-change features, 17 showed nominally significant treatment-adjusted correlations (P < 0.05), but none remained significant after BH correction (q < 0.10; **Figure 5E; Table S8**). Together, these findings indicate that associations between *Bacteroides* abundance and individual tumor, serum, and fecal metabolites, as well as between tumor metabolic modules and tumor burden, were largely attenuated after adjustment for treatment-group effects.

**Figure 5.**
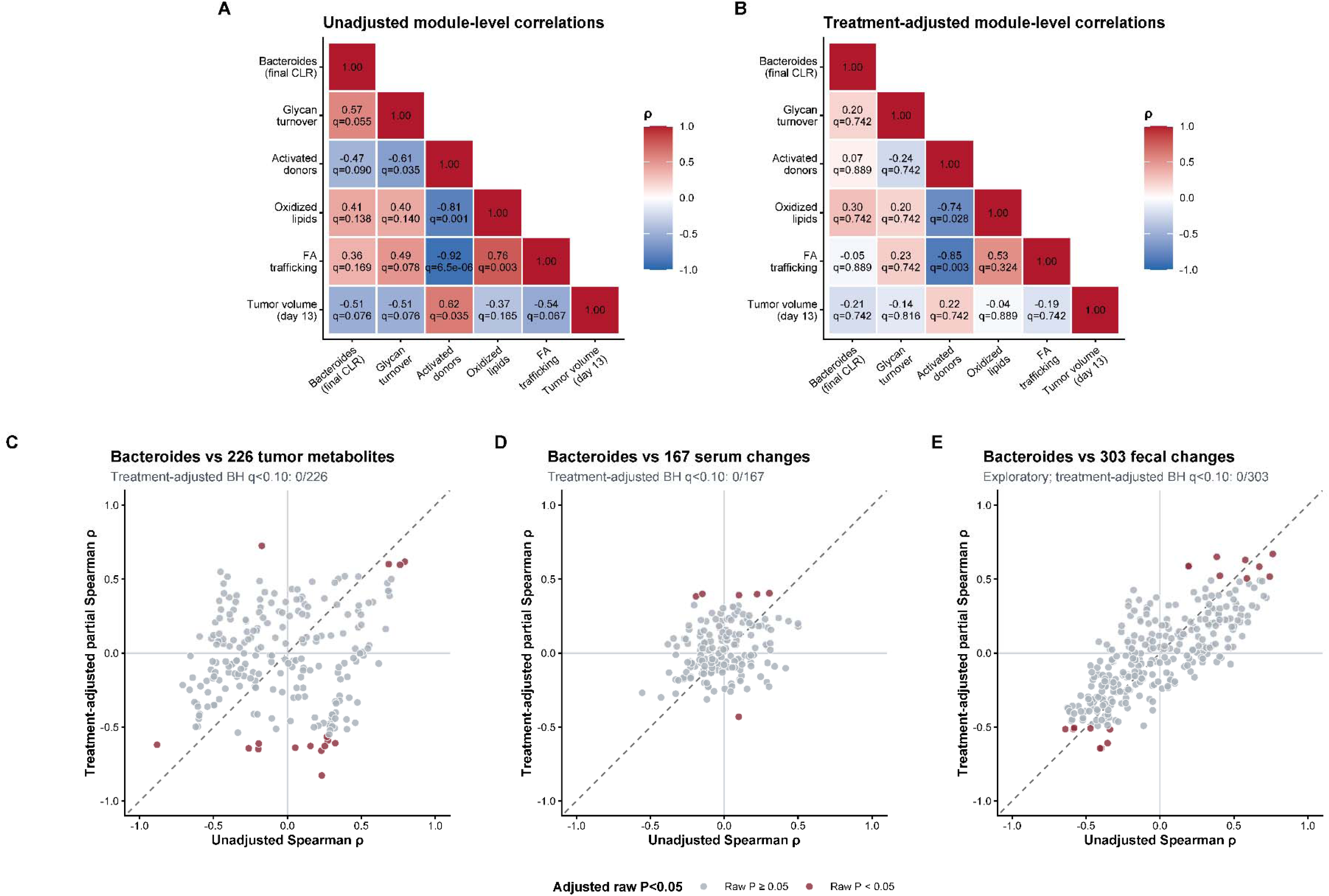
Treatment adjustment of cross-omics associations. Matched mouse-level analyses compare unadjusted Spearman correlations with treatment-adjusted partial Spearman correlations obtained after rank residualization on BO, α-mPD-1, and BO × α-mPD-1. **(A,B)** Lower-triangle module-level matrices for final Bacteroides CLR, four tumor-metabolic module scores, and day-14 tumor volume (n=16); cells display ρ and BH q, and color encodes ρ**. (C–E)** Feature-wide Bacteroides association screens for 226 tumor metabolites, 167 serum paired-change metabolites, and 303 fecal paired-change metabolites, respectively. Points highlighted in the feature-wide panels have treatment-adjusted raw P<0.05. The fecal analysis is exploratory because of the smaller, unbalanced paired cohort; its subtitle reports treatment-adjusted BH q<0.10 for 0/303 features. Dashed diagonals indicate unchanged ρ after treatment adjustment. BH correction is performed separately within each omics block and correlation method.

### 3.7 Cross-omics integration identified coordinated microbial and metabolic responses

To integrate microbiome composition, fecal and serum metabolomics, tumor metabolites, and tumor burden, we constructed a targeted cross-omics correlation network comprising 25 nodes selected independently of network correlation P values (**Figure 6; Table S9-1; Table S9-2**). The network included six microbial genera, five fecal metabolites, five serum metabolites, eight tumor metabolites representing four predefined metabolic modules, and day-14 tumor volume. Of 237 possible cross-block feature pairs, 197 had at least 16 matched observations and were included in the correlation analysis. The unadjusted network contained 42 associations at the nominal P < 0.05 threshold, of which 11 remained significant after network-wide Benjamini–Hochberg correction (q < 0.05; **Figure 6A; Table S9-1; Table S9-2**).

**Figure 6.**
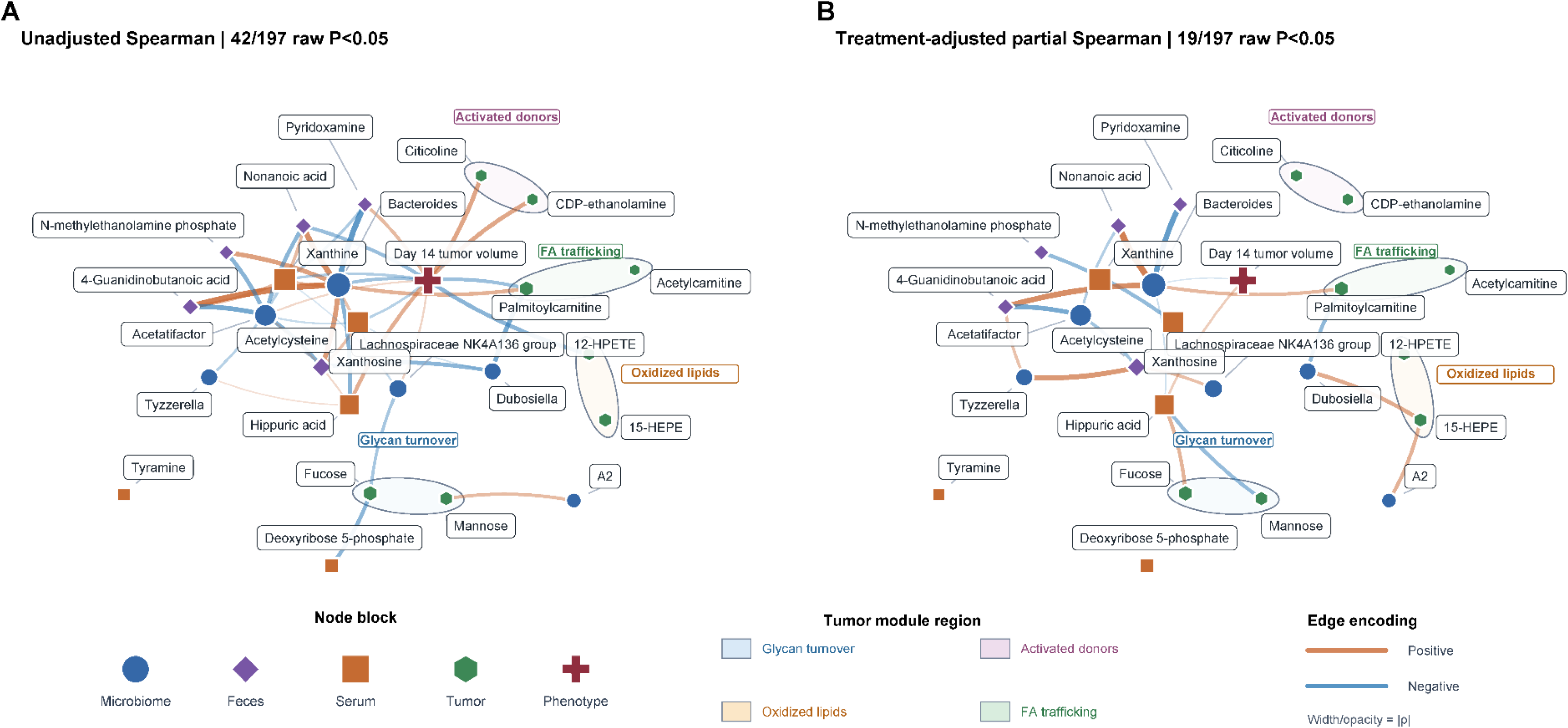
Integrated cross-omics correlation network before and after treatment adjustment. The network contains 25 nodes: 6 genus-level microbiome features (circles), 5 fecal metabolites (diamonds), 5 serum metabolites (squares), 8 tumor metabolites (hexagons), and day-14 tumor volume (cross). The panels are arranged side by side with identical saved node coordinates: **(A, left)** unadjusted Spearman network; **(B, right)** treatment-adjusted partial Spearman network after rank residualization on BO, α-mPD-1, and their interaction. Only cross-block pairs were tested. Of 237 candidate pairs, 197 met the pairwise eligibility rule and 40 were excluded. Panels display raw P<0.05 edges (42 unadjusted; 19 adjusted), a visualization threshold rather than an FDR threshold. Edge color indicates sign; width and opacity encode |ρ|. Weaker edges are drawn before stronger edges. Node size is degree calculated from the union of displayed edges.

*Bacteroides* was among the most highly connected nodes in the unadjusted network, with 10 associations meeting the nominal P < 0.05 threshold. Greater baseline-to-final increases in *Bacteroides* CLR abundance were positively correlated with fecal nonanoic acid, 4-guanidinobutanoic acid, acetylcysteine, and N-methylethanolamine phosphate, and negatively correlated with fecal pyridoxamine. *Bacteroides* was also positively associated with serum xanthine and xanthosine and negatively associated with serum hippuric acid and day-14 tumor volume. Conversely, serum hippuric acid was positively correlated with day-14 tumor volume (ρ = 0.577, q = 0.0094), suggesting distinct relationships between microbiome-associated metabolic features and tumor burden before treatment adjustment (**Figure 6A; Table S9-1**). After adjusting for BO, α-mPD-1, and their interaction, the number of nominally significant network edges decreased from 42 to 19, while the number surviving network-wide BH correction decreased from 11 to one (**Figure 6B; Table S9-1**). The only association that remained significant after FDR correction was the positive correlation between *Bacteroides* paired CLR change and fecal nonanoic acid paired metabolic change (partial ρ = 0.798, q = 0.041; n = 19). Associations of *Bacteroides* with fecal 4-guanidinobutanoic acid, fecal pyridoxamine, serum hippuric acid, and day-14 tumor volume remained nominally significant (P < 0.05) but did not survive FDR correction. Together, these results suggest that many of the observed cross-omics associations were influenced by treatment-group differences, whereas the relationship between *Bacteroides* and fecal nonanoic acid remained significant after accounting for treatment assignment.

## 4. Discussion

In this study, BO administration induced moderate tumor-growth suppression, whereas the BO + α-mPD-1 group showed the lowest tumor burden across the longitudinal analysis (**Figure 7**). However, the BO × α-mPD-1 interaction was not significant on day-14 using the linear regression model; therefore, the present data supports a strong combination-associated phenotype but does not establish statistical synergy. These findings extend a substantial body of work demonstrating that the gut microbiome can influence responsiveness to PD-1 blockade in both preclinical models and patients.[4,6,7] More broadly, microbiome-targeted interventions have emerged as a potential strategy for modifying immune checkpoint-blockade responsiveness, although effects vary substantially depends on microbial composition, host context, and tumor type.[28]

**Figure 7.**
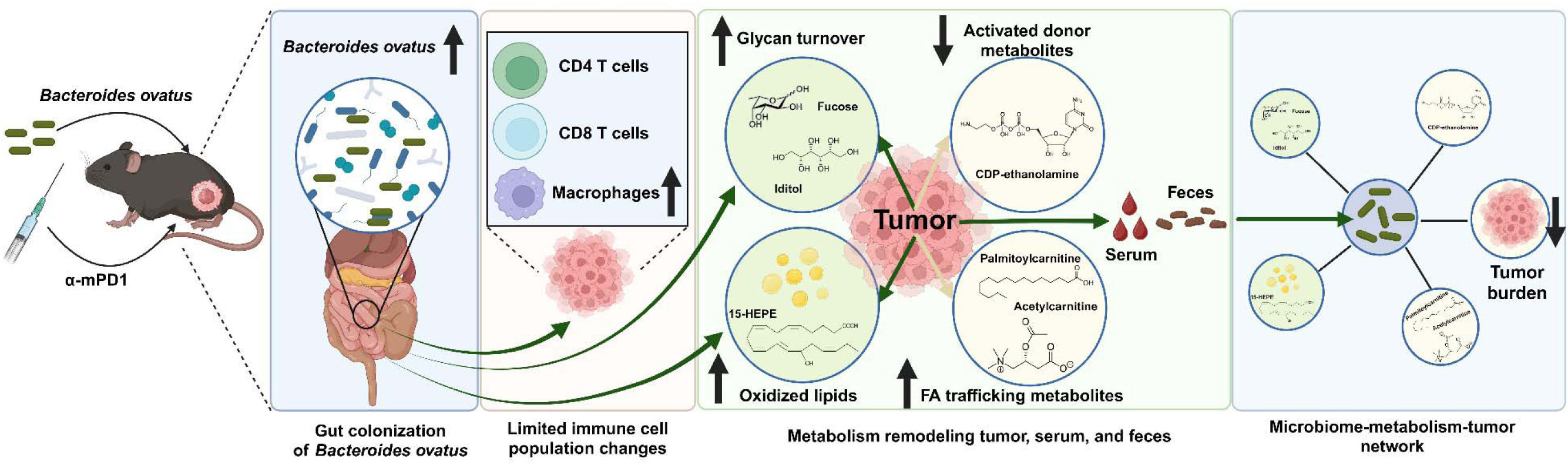
Working model of *Bacteroides ovatus*–associated microbiome and metabolic remodeling during α-mPD-1 therapy in the MC38 colorectal cancer model. Oral administration of *Bacteroides ovatus* (BO) increased fecal Bacteroides abundance and was associated with broader remodeling of the gut microbial community. Exploratory tumor immune profiling showed relatively limited changes in the measured T-cell populations, with a modest increase in macrophage abundance in the combination group. In contrast, tumor metabolomics identified pronounced treatment-associated remodeling characterized by increased glycan turnover, oxidized lipids, and fatty acid trafficking modules together with decreased activated-donor metabolites in the BO + α-mPD-1 group. Longitudinal serum and fecal metabolomics further indicated systemic and intestinal metabolic changes, although the fecal analysis was exploratory. Integrated cross-omics analyses linked microbial and metabolic features with tumor burden, but many associations were attenuated after adjustment for treatment assignment. The schematic summarizes the observed microbiome–metabolism–tumor relationships and represents a hypothesis-generating model rather than a demonstrated causal pathway.

BO administration produced a marked increase in *Bacteroides* together with broader changes in microbial community structure (**Figure 7**), including changes that remained detectable after *Bacteroides* itself was removed from the ordination analyses. This indicates that BO colonization altered the surrounding microbial ecosystem rather than simply increasing a single genus. The tumor qPCR data were also suggestive of increased BO-associated DNA signal in BO-exposed tumor culture samples, although these measurements may need further validation and cannot capture viable bacterial translocation or intratumoral colonization. Experimental evidence from other checkpoint-blockade models has shown that gut bacteria can reach extraintestinal sites and contribute to antitumor immunity, providing biological precedent for distant microbiome–tumor interactions.[29,30] Nevertheless, culture, species-resolved sequencing, and spatial localization will be required to determine whether BO itself reaches and persists within the tumor in the future studies.

Tumor metabolomics revealed coordinated remodeling of glycan turnover, activated-donor metabolism, oxidized lipids, and fatty-acid trafficking (**Figure 7**). These changes were most pronounced in the BO enhanced anti-PD1 group and provide a metabolic context for the observed tumor phenotype. Importantly, static metabolite abundance cannot establish pathway flux or identify whether individual metabolites originate from microbial, tumor, stromal, or immune compartments. Nevertheless, the findings are consistent with increasing evidence that microbial metabolites and host metabolic reprograming can influence antitumor immunity and checkpoint-blockade response.[31,32] Recent work has also reported metabolite-dependent antitumor effects of *B. ovatus* in colorectal cancer, supporting the broader concept that this species can influence tumor biology through metabolic mechanisms, although the specific metabolites and pathways differ from those identified here.[11] Longitudinal serum and fecal metabolomics further suggested that these metabolic effects extend beyond the tumor, although the strength of evidence differed by compartment (**Figure 7**). Serum paired-change profiles showed treatment-associated structure, whereas fecal metabolomics was less definitive and remained exploratory because of its limited sample availability. Together, these observations suggest that BO-associated metabolic remodeling may involve both local and systemic compartments (**Figure 7**).

Meanwhile, treatment adjustment substantially changed the cross-omics interpretation. Many associations among *Bacteroides*, metabolic features, and Day-14 tumor volume were attenuated after accounting for BO, α-mPD-1, and their interaction, indicating that much of the unadjusted covariance reflected treatment-group separation. Similarly, the integrated network contracted from 42 to 19 raw-P<0.05 edges after treatment adjustment, and only the *Bacteroides*–nonanoic acid association remained significant after network-wide BH correction. These results argue against interpreting cross-group correlations as evidence of mediation. Instead, the adjusted network identifies a smaller set of relationships that warrant direct mechanistic testing, particularly microbiome–metabolite associations that persist after removal of treatment-associated structure.

Exploratory immune profiling did not reveal broad changes across the measured T-cell activation, exhaustion, and memory populations, although interpretation is limited by the small number of animals. Thus, the present data cannot exclude immune mechanisms that involve functional, transcriptional, spatial, or metabolic states rather than changes in cell frequency. This distinction is particularly relevant because tumor metabolism can strongly influence immune-cell function even in the absence of major changes in immune-cell abundance.

Several limitations should therefore be acknowledged. First of all, the MC38 model does not capture the heterogeneity of human CRC, the tumor-metabolomics cohort was modest, and fecal metabolomics was exploratory. Additionally, the current multi-omics analyses provided a strong association network and will enable additional hypothesis testing studies in the future, but our current results do not establish mediation or causality. Future studies should combine species-resolved microbiome profiling with independently replicated tumor localization, targeted metabolite quantification, stable-isotope tracing, and metabolite perturbation experiments. Such studies will be necessary to determine whether the metabolic relationships identified here are mechanistically involved in the tumor response to BO and PD-1 blockade.

## 5. Conclusion

In conclusion, BO administration was associated with substantial remodeling of the gut microbiome and tumor metabolic landscape during α-mPD-1 treatment, while the BO + α-mPD-1 group exhibited the lowest tumor burden in this CRC model. Although the study did not establish a statistically significant BO × α-mPD-1 interaction or a causal metabolic mechanism, the integrated analyses identified coordinated changes in tumor metabolic pathways and a smaller subset of microbiome–metabolite associations that persisted after adjustment for treatment assignment. These findings support a model in which BO-associated ecological and metabolic changes accompany improved tumor control during PD-1 blockade and provide specific microbial–metabolic relationships for future mechanistic validation. Further studies using larger cohorts, species-resolved microbiome profiling, targeted metabolite quantification, spatial localization, and functional perturbation will be required to determine whether these associations directly contribute to immunotherapy response.

## Appendices

Appendices for supplementary materials include **Figure S1-S7 and Table S1-S9** All tables are attached separately.

## Data availability statement

All raw data for the metabolomics have been deposited to MassIVE under accession number MSV000103891.

## Acknowledgement

We thank Dr. Daniel Spakowicz for gifting the MC38 cell lines used in this study.

## Disclosure of generative AI use

During the preparation of this manuscript, the authors used OpenAI Codex (GPT-6 Sol Ultra) to assist in developing statistical analysis workflows, recommending statistical approaches, and generating R scripts for data processing, statistical analyses, and data visualization based on the original experimental datasets and research objectives. The AI-generated scripts were subsequently executed by the authors in R to perform the analyses and generate the corresponding figures. Additionally, ChatGPT (OpenAI, GPT-6) was used to assist with English language editing, grammatical corrections, and improving the clarity and readability of the manuscript. The authors retained full responsibility for evaluating the appropriateness of the statistical methods, reviewing and verifying the analytical code and results, interpreting the findings, and ensuring the scientific integrity of the manuscript. The authors confirm the originality and accuracy of the final manuscript, have reviewed the applicable terms of use of the AI tools and their suitability for publication, and take full responsibility for all analyses, results, figures, references, and the final published content.

## CRediT Roles

H.Z., S.Z., and S.G. contributed to the mouse experiments and collected samples and data for metabolomics, microbiome, and FACS analyses. H.Z., S.G., H.C. prepared samples and H.C. analyzed the FACS data. F.X. contributed to microbiome data collection and sample preparation. S.G., X.P., H.J., C.G., A.G., O.L., and N.N. contributed to sample collection. F.C. contributed to MC38 cell culturing for the subcutaneous model construction. H.Z. contributed to data analysis, manuscript preparation, and figure generation. H.S., J.L., and J.Z. conceived the project, acquired funding, and revised the manuscript.

---

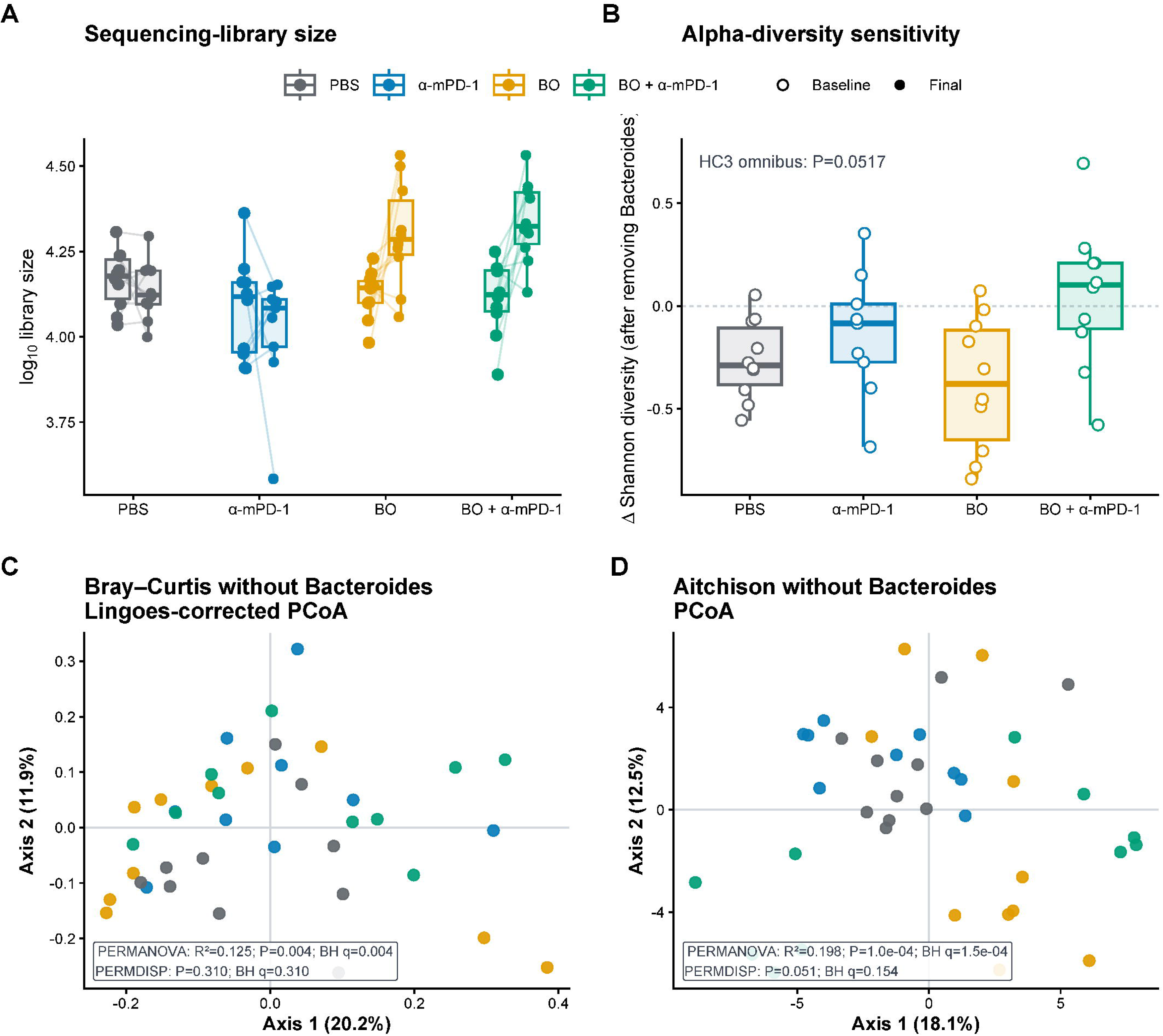

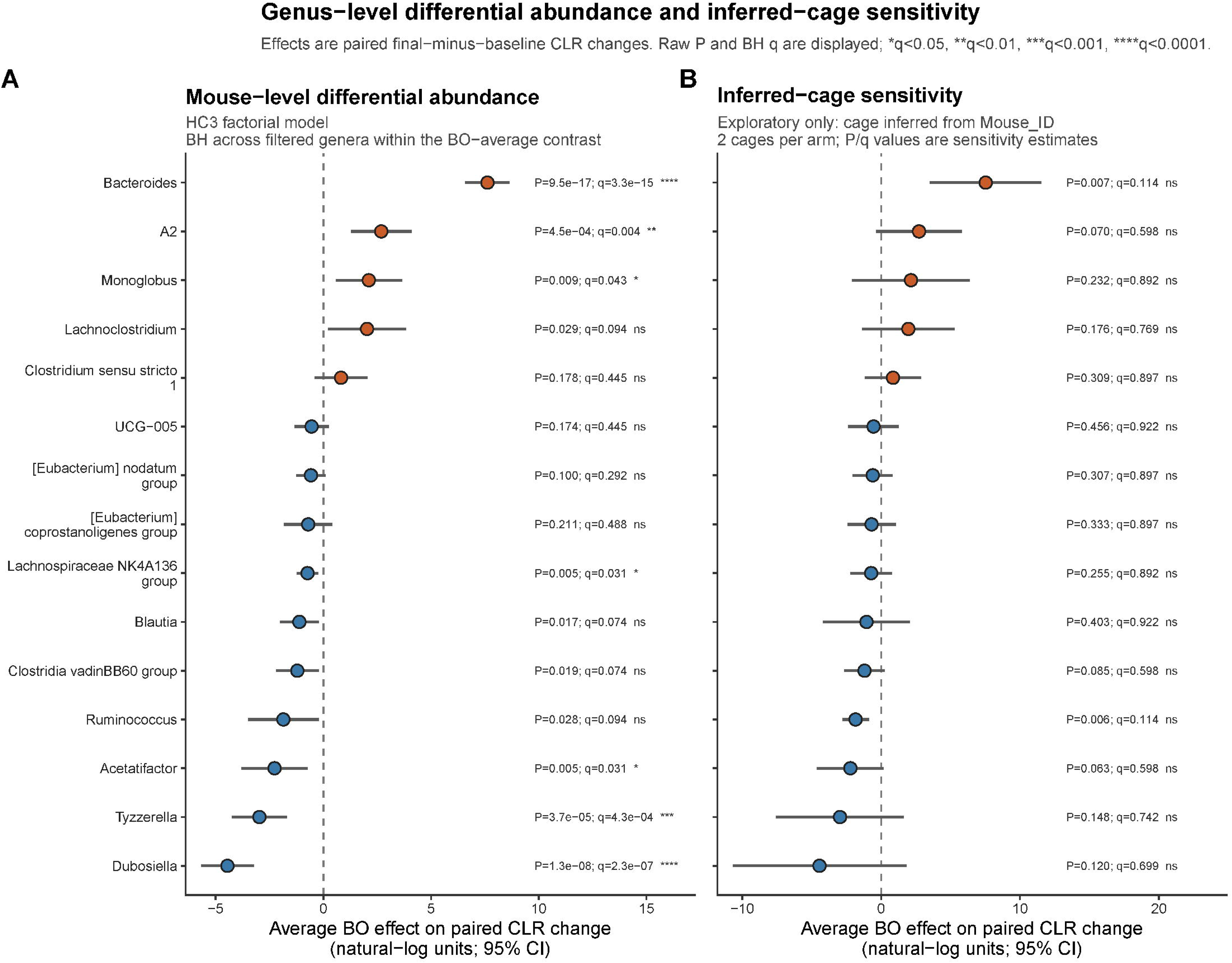

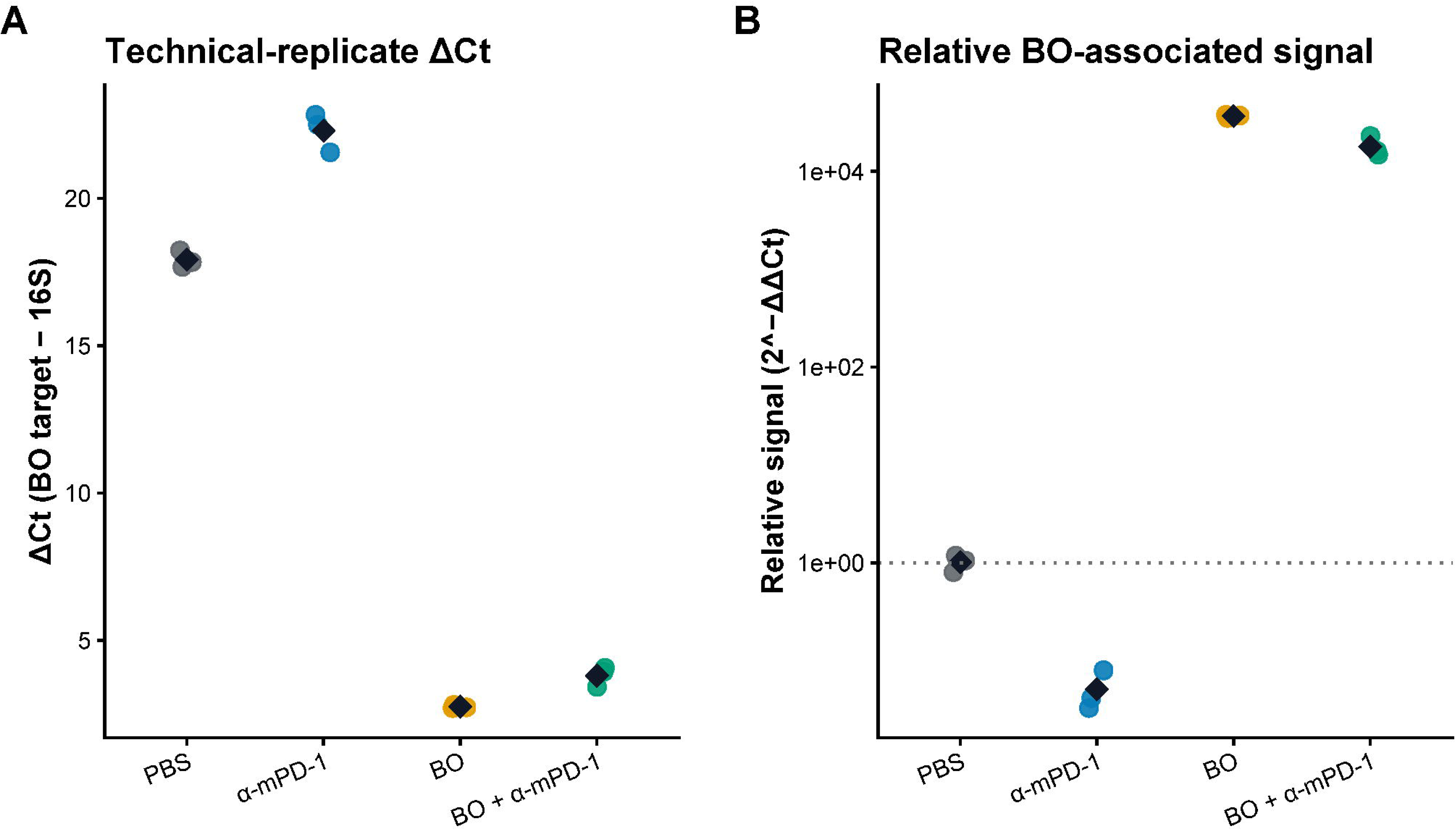

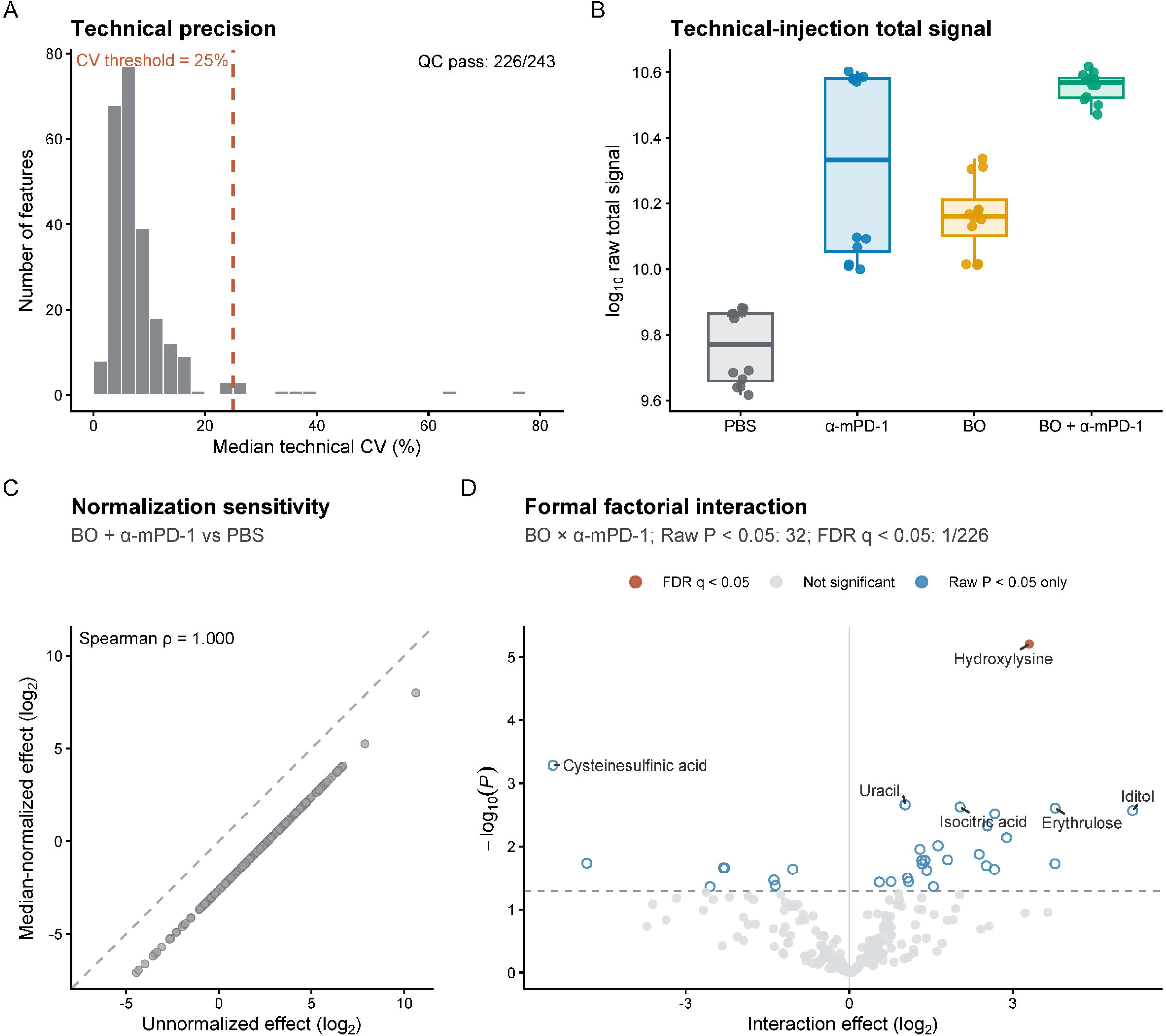

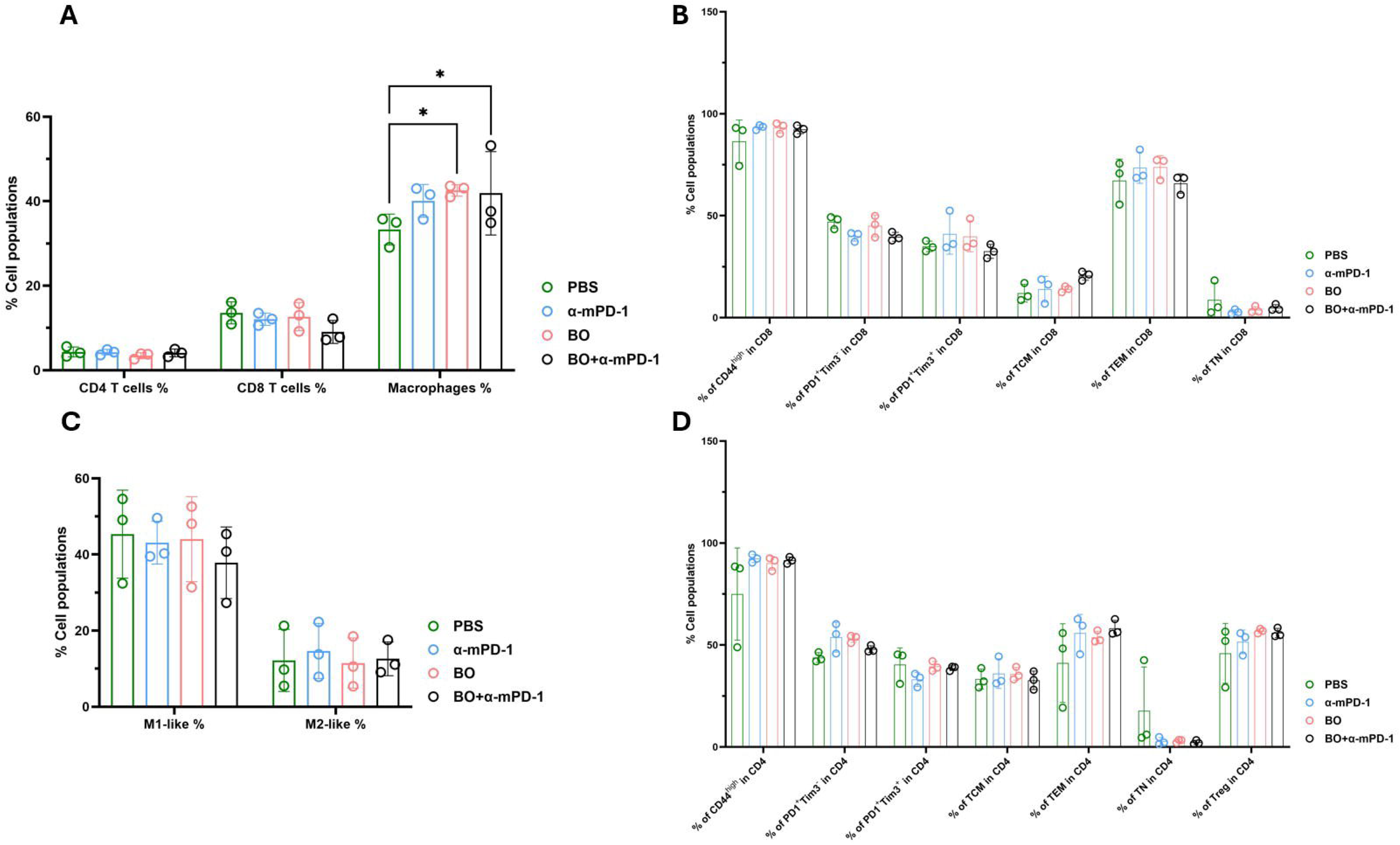

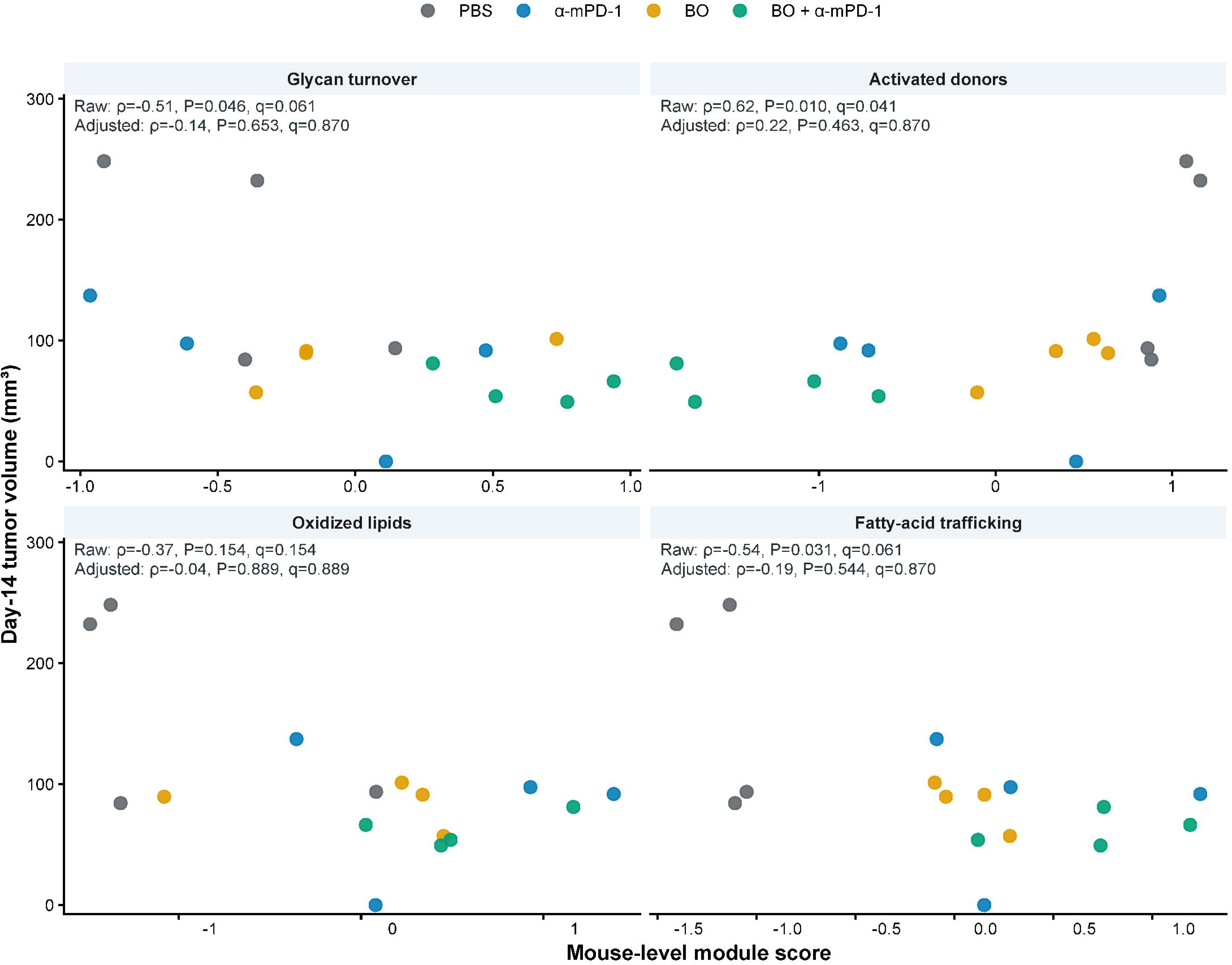

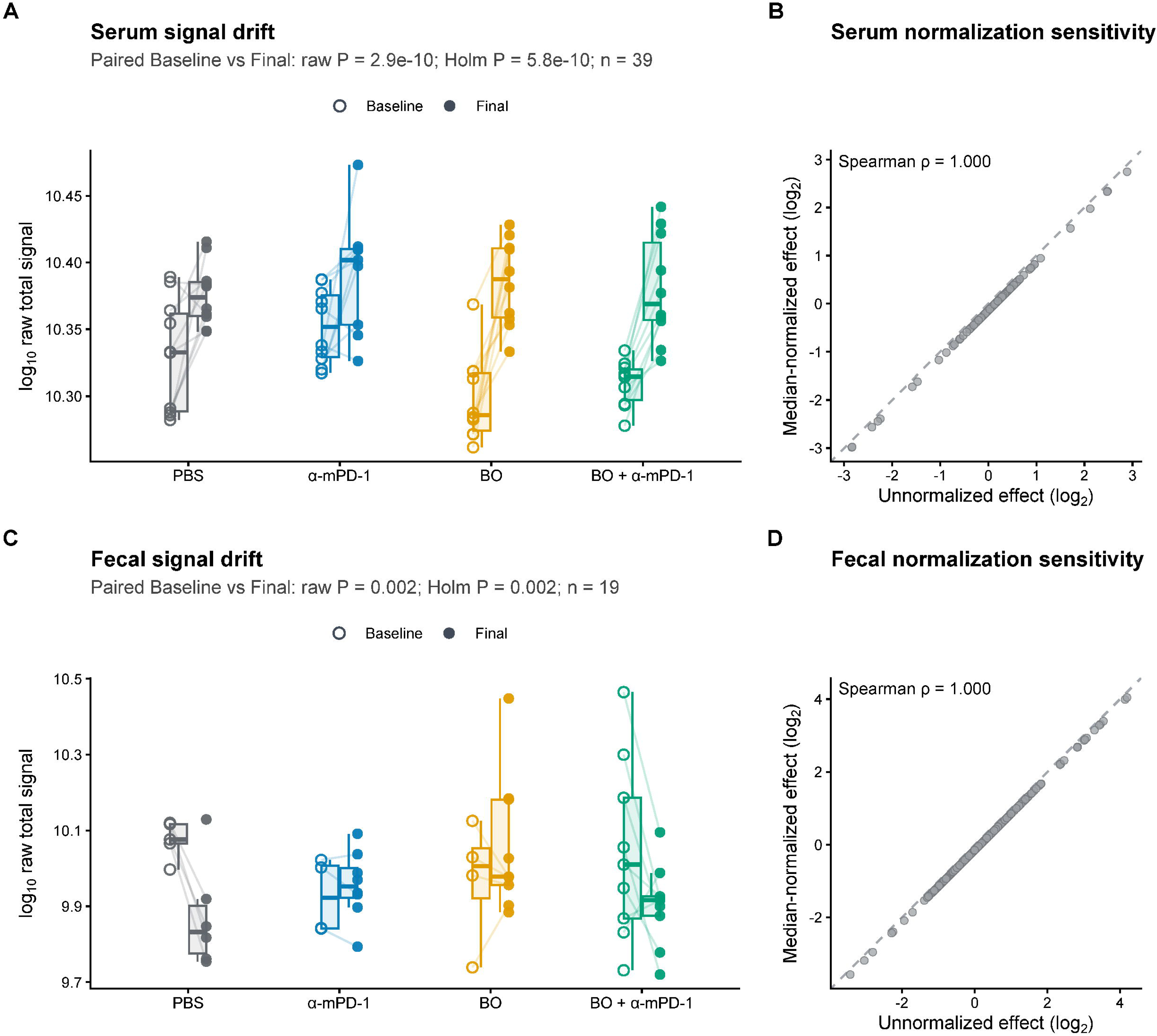

