## supplementary materials for "*Bacteroides ovatus* administration enhanced α-mPD-1 immunotherapy in a mouse colorectal cancer model"

Huan Zhang^1, 2, 5^, Shiqi Zhang^1, 2, 5^, Heng-Yi Chen^2, 3^, Mengyang Xu^4^, Shuaixin Gao^1,2^, Xinru Pang^1, 2^, Li Chen^1, 2^, Hyein Jung^1, 2^, Chao Guo^1, 2^, Andrew Gold^1, 2^, Natalie Crawford^1, 2^, Fouad Choueiry^1, 2^, Owen Laib^1, 2^, Haifei Shi^4^, Chan-Wang Jerry Lio^2, 3^, Jiangjiang Zhu^1, 2*^

**Author affiliation:**

1. Human Nutrition Program, Department of Human Sciences, The Ohio State University, Columbus, OH, 43210, USA.
2. James Comprehensive Cancer Center, The Ohio State University, Columbus, OH, 43210, USA.
3. Department of Microbial Infection and Immunity, The Ohio State University, Columbus, OH, 43210, USA.
4. Department of Biology, Miami University, Oxford, OH, 45056, USA.
5. These authors contributed to the work equally.

Corresponding author:

*Jiangjiang Zhu, Ph.D., Comprehensive Cancer Center and Department of Human Sciences, The Ohio State University, 400 W 12th Ave, Columbus, OH 43210 Tel: 614-685-2226,

**Supplementary figure legends:**

**Figure S1. Microbiome quality control and sensitivity analyses.** Paired baseline/final microbiome samples from 39 mice. **(A)** log_10_ sequencing-library size by group and time; open points are baseline, filled points final, and lines connect mice. Boxes show median and IQR with 1.5×IQR whiskers. **(B)** Final-minus-baseline Shannon change after removing *Bacteroides*; the HC3 four-group omnibus raw P is shown. **(C)** Final Bray–Curtis PCoA after Bacteroides removal, with Lingoes correction. **(D)** Final Aitchison PCoA after Bacteroides removal; a 0.5-read pseudocount and CLR transformation were used after prevalence/abundance filtering. The locked Bray–Curtis-without-Bacteroides tests were PERMANOVA R²=0.1247; raw P=0.0037; BH q=0.0037; PERMDISP raw P=0.3098; BH q=0.3098; the locked Aitchison-without-Bacteroides tests were PERMANOVA R²=0.1979; raw P=0.0001; BH q=0.00015; PERMDISP raw P=0.0515; BH q=0.1545. Colors identify PBS, α-mPD-1, BO, and BO + α-mPD-1.

**Figure S2. Genus-level differential abundance and inferred-cage sensitivity.** Differential abundance was evaluated on within-mouse final-minus-baseline CLR changes for 35 retained genera across 39 mice. **(A)** Mouse-level average BO effect from 2 × 2 HC3 models. Points are effect estimates, bars are 95% CIs, and the text gives exact raw P and BH-adjusted q values within the 35-genus BO-effect family. Asterisk labels reflect BH-adjusted q values: ns, q≥0.05; q<0.05; **q<0.01; ***q<0.001; ****q<0.0001. **(B)** Descriptive cage-mean sensitivity for the same displayed genera using eight inferred cages (two per arm). Points and intervals are cage-level effect estimates and 95% CIs. Because cage identity was inferred from mouse IDs and only two cages were available per arm, Panel B is not treated as randomized cage-level inference.

**Figure S3. BO-specific qPCR signal in anaerobically expanded tumor-derived bacterial cultures. (A)** ΔCt values calculated as Cq(BO target) − Cq(16S); lower ΔCt values indicate a greater BO-associated signal relative to total bacterial 16S signal. **(B)** Relative BO-associated signal calculated using the 2^−ΔΔCt^ method, where ΔΔCt = ΔCt(sample) − mean ΔCt(PBS). Values are shown on a logarithmic scale, with the PBS group serving as the reference (relative signal = 1). Circles represent individual technical qPCR replicates and diamonds indicate group arithmetic means.

**Figure S4. Tumor-metabolomics quality control and interaction sensitivity.** Tumor metabolomics comprised 16 biological mice, four per group, with three technical injections per mouse. **(A)** Distribution of feature-level median technical CV before aggregation; the dashed line marks the 25% median-CV threshold. The full rule also required no more than 25% of mice to have CV>30%, yielding 226/243 QC-passing features. **(B)** Technical-injection total signal, showing raw total signal for each technical injection by group; boxes show median and IQR, whiskers 1.5×IQR, and points are technical injections and are not inferential units. **(C)** Comparison of BO + α-mPD-1 versus PBS effect estimates from unnormalized log₂ and median-normalized log₂ HC3 models; each point is a QC-passing feature and the dashed line is equality. **(D)** Volcano plot for the formal BO × α-mPD-1 interaction. Filled red points meet BH q<0.05, open blue points raw P<0.05 only, and gray points neither; 32/226 had raw P<0.05 and 1/226 retained BH q<0.05.

**Figure S5. Flow cytometric analysis of immune cell populations across the PBS, α-mPD-1, B. ovatus (BO), and BO+α-mPD-1 treatment groups. (A)** shows the percentages of CD4 T cells, CD8 T cells, and macrophages among total cell populations, with significant differences in macrophage abundance indicated between the marked groups. **(B)** shows the distribution of CD8 T-cell subsets, including CD44^hi^, PD1^+^Tim3^−^, PD1^+^Tim3^+^, central memory T (TCM), effector memory T (TEM), and naïve T (TN) cells. Panel **(C)** shows the proportions of M1-like and M2-like macrophages. Panel **(D)** shows the distribution of CD4 T-cell subsets, including CD44^high^, PD1^+^Tim3^−^, PD1^+^Tim3^+^, TCM, TEM, TN, and regulatory T (Treg) cells. Data are presented as mean ± SD, and each symbol represents one biological replicate. P < 0.05 for the indicated comparisons*.*

**Figure S6. Tumor metabolic modules and day-14 tumor burden before and after treatment adjustment.** Scatter plots show day-14 tumor volume versus mouse-level module score in the 16-mouse tumor-metabolomics cohort: **(A)** glycan turnover, **(B)** activated donors, **(C)** oxidized lipids, and **(D)** fatty-acid trafficking. Points are biological mice colored by treatment. Each panel reports unadjusted Spearman rho, raw P, and BH q across the four-module family, followed by treatment-adjusted partial Spearman rho, P, and q after rank residualization on BO, α-mPD-1, and BO × α-mPD-1.

**Figure S7. Longitudinal serum and fecal metabolomics signal and normalization sensitivity.** **(A)** Serum log_10_ raw total signal at baseline and final time points for 39 paired mice. **(B)** Comparison of feature-effect estimates from unnormalized and median-normalized serum paired-change models. **(C)** Fecal log₁₀ raw total signal for 19 paired mice. **(D)** Equivalent fecal normalization comparison. In A and C, open points are baseline, filled points final, lines connect mice, and boxes show median/IQR with 1.5×IQR whiskers. Overall baseline-versus-final signal tests used paired comparisons and Holm correction across the serum/fecal diagnostic family. In B and D, each point is a feature, the dashed line is equality, and Spearman rho summarizes effect-ranking concordance. These panels diagnose global signal drift and do not replace the normalized feature-wise inference.

**Figure S1**

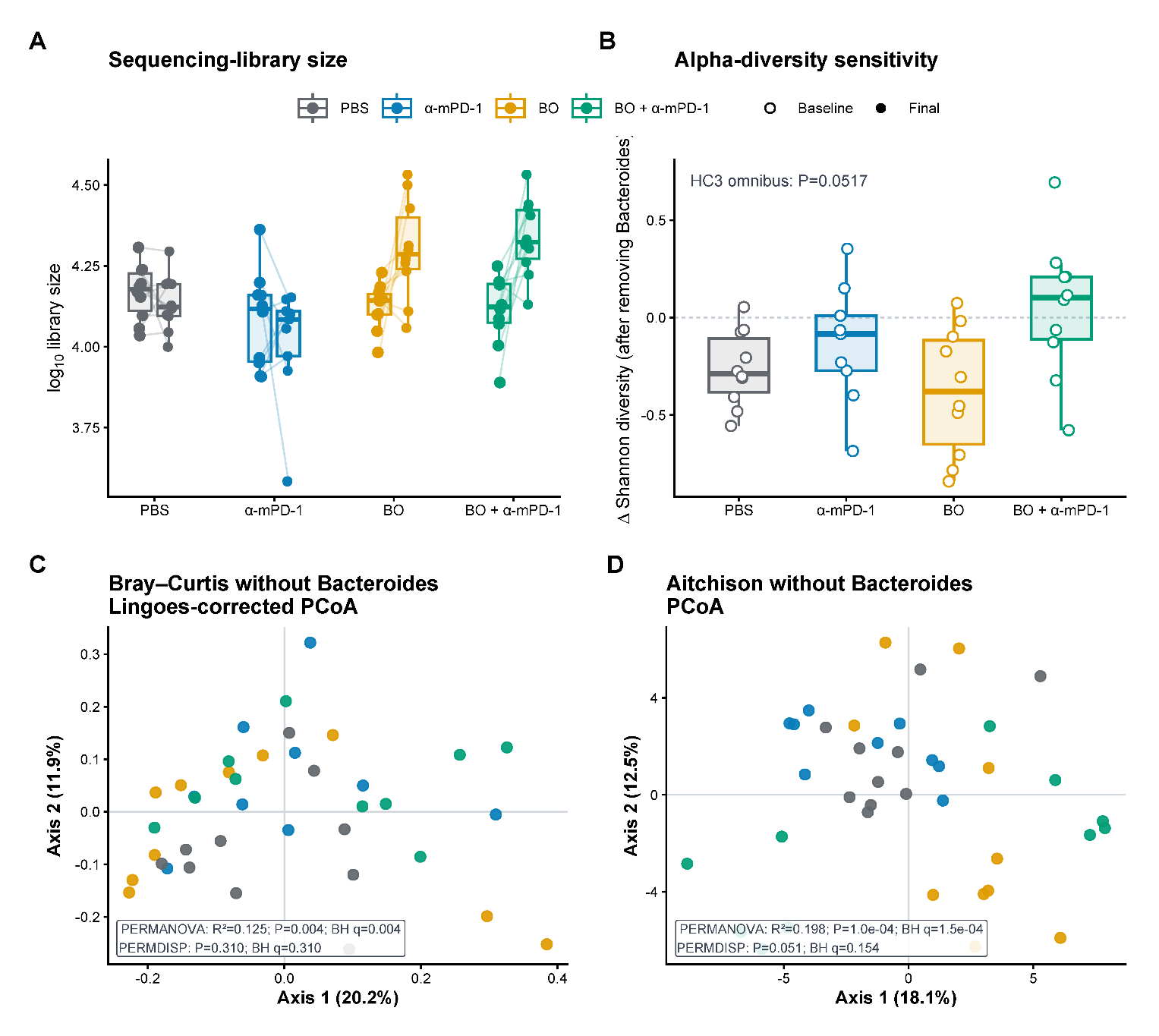

**Figure S2**

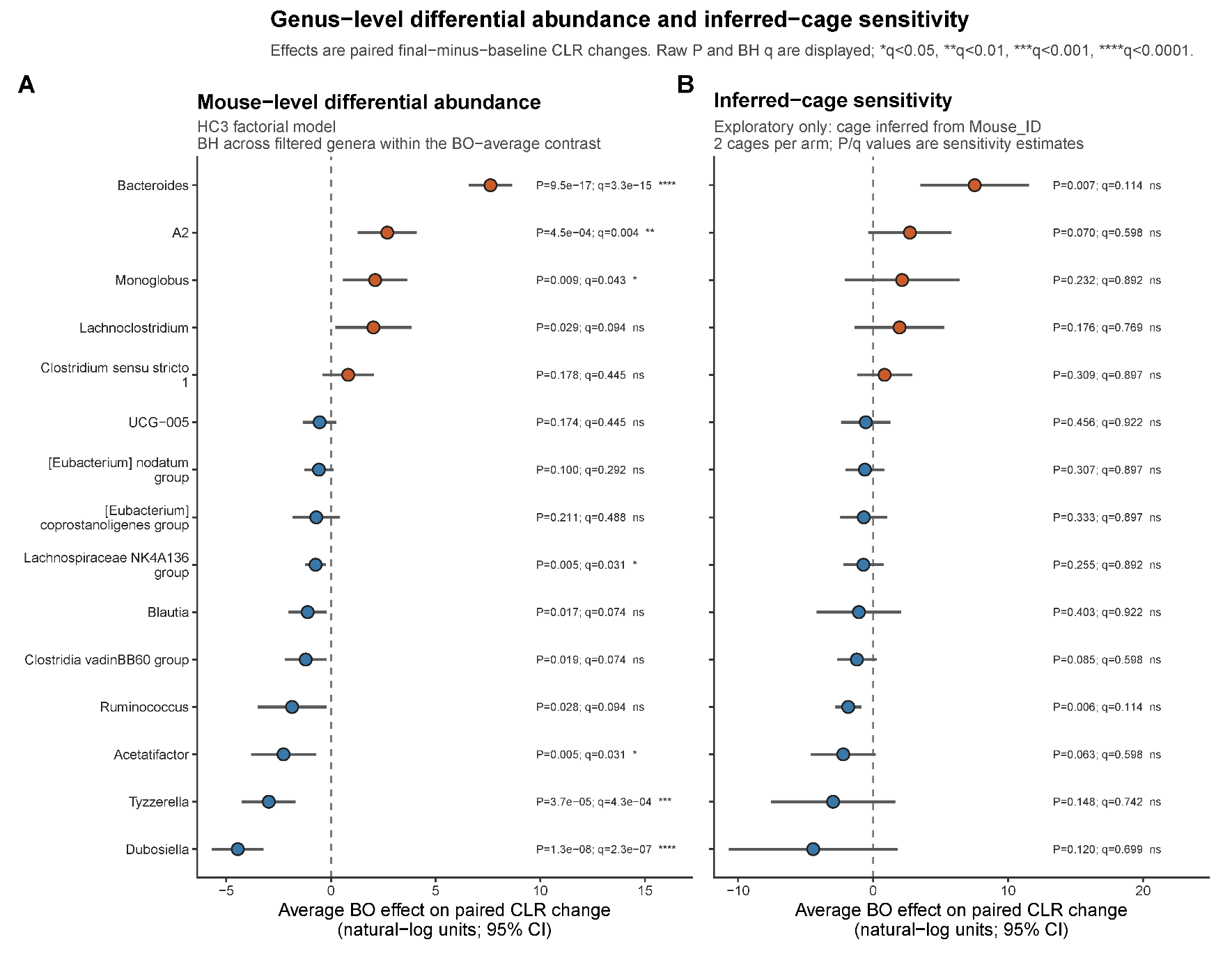

**Figure S3**

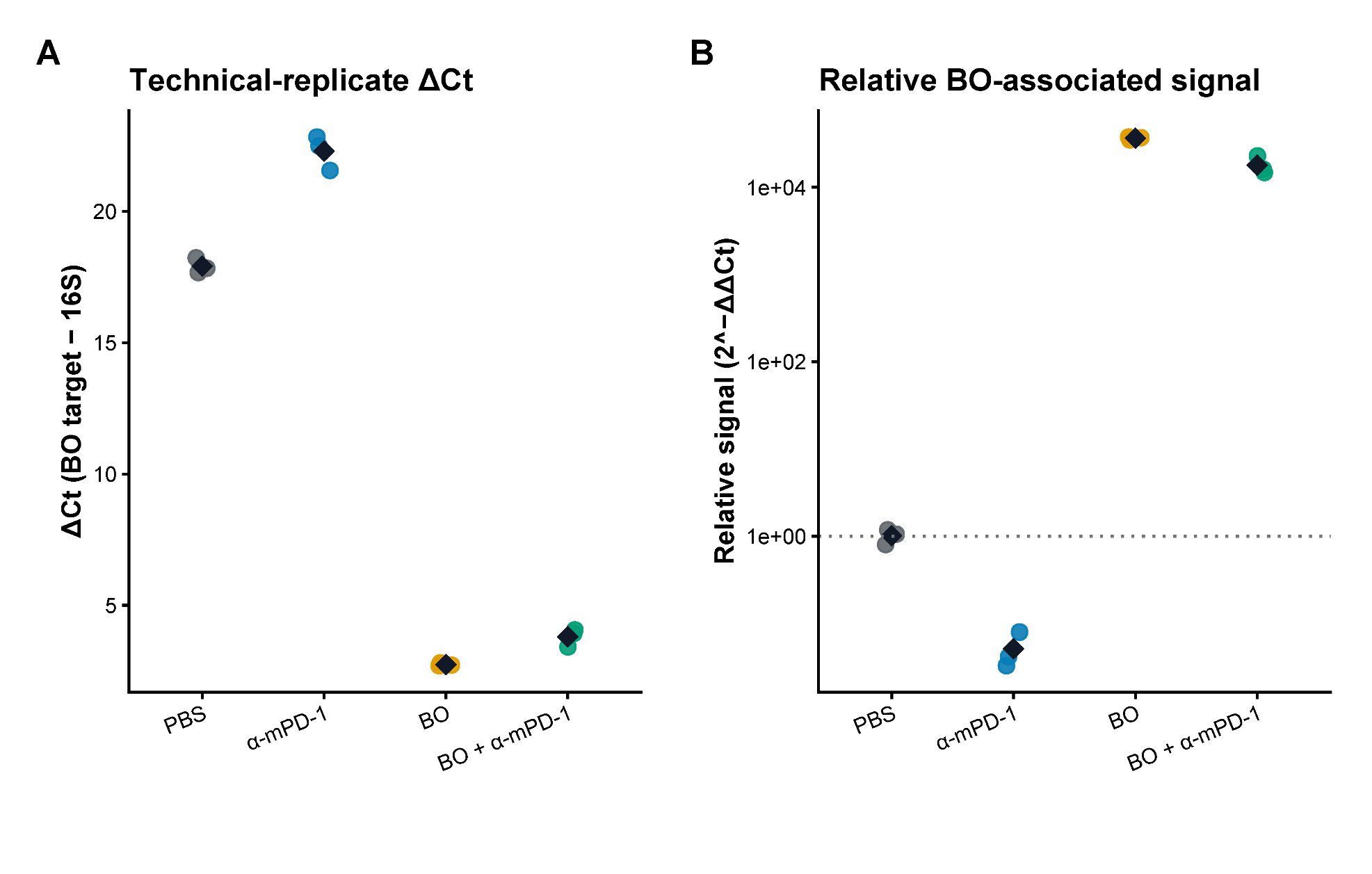

**Figure S4**

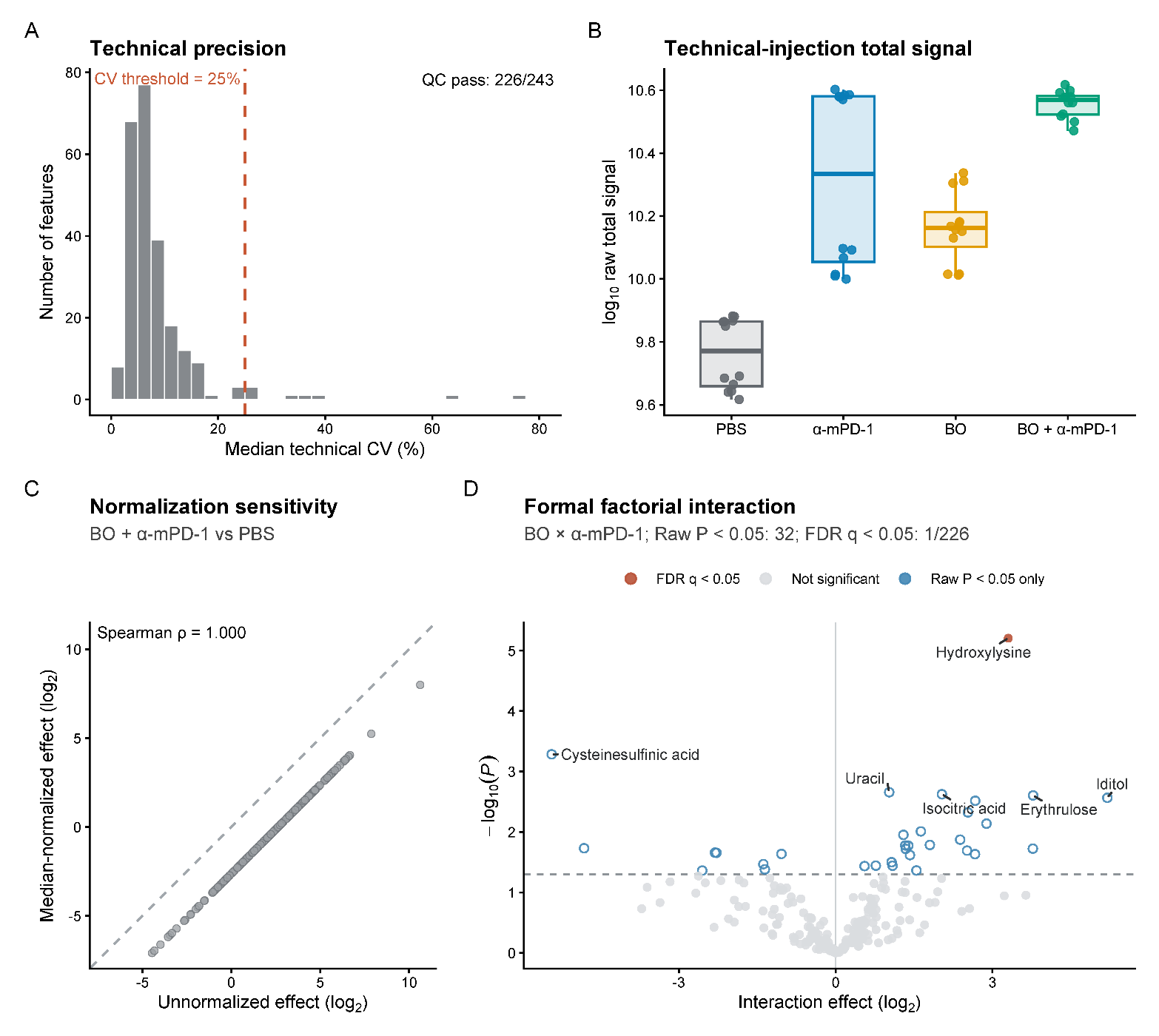

**Figure S5**

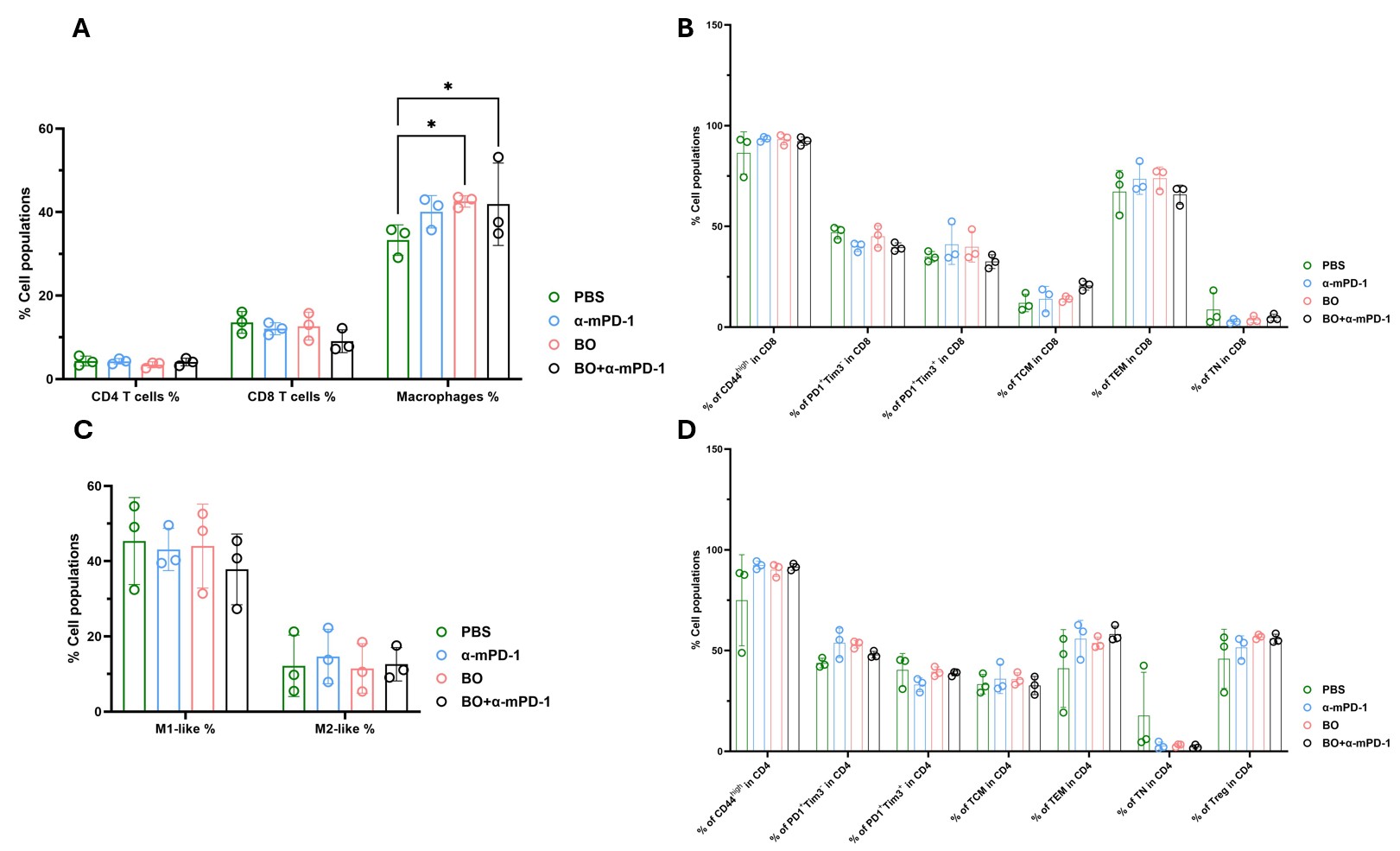

**Figure S6**

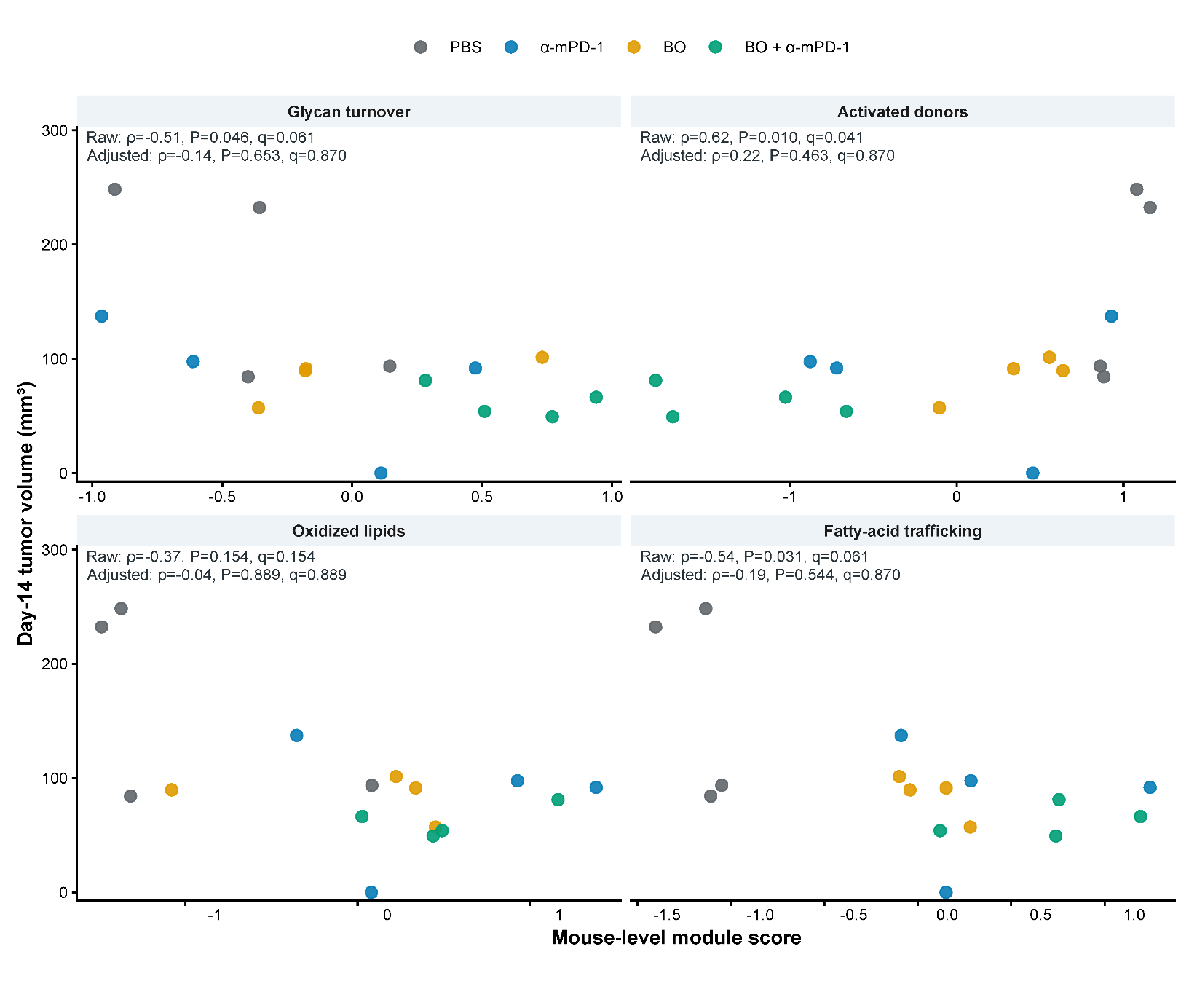

**Figure S7**

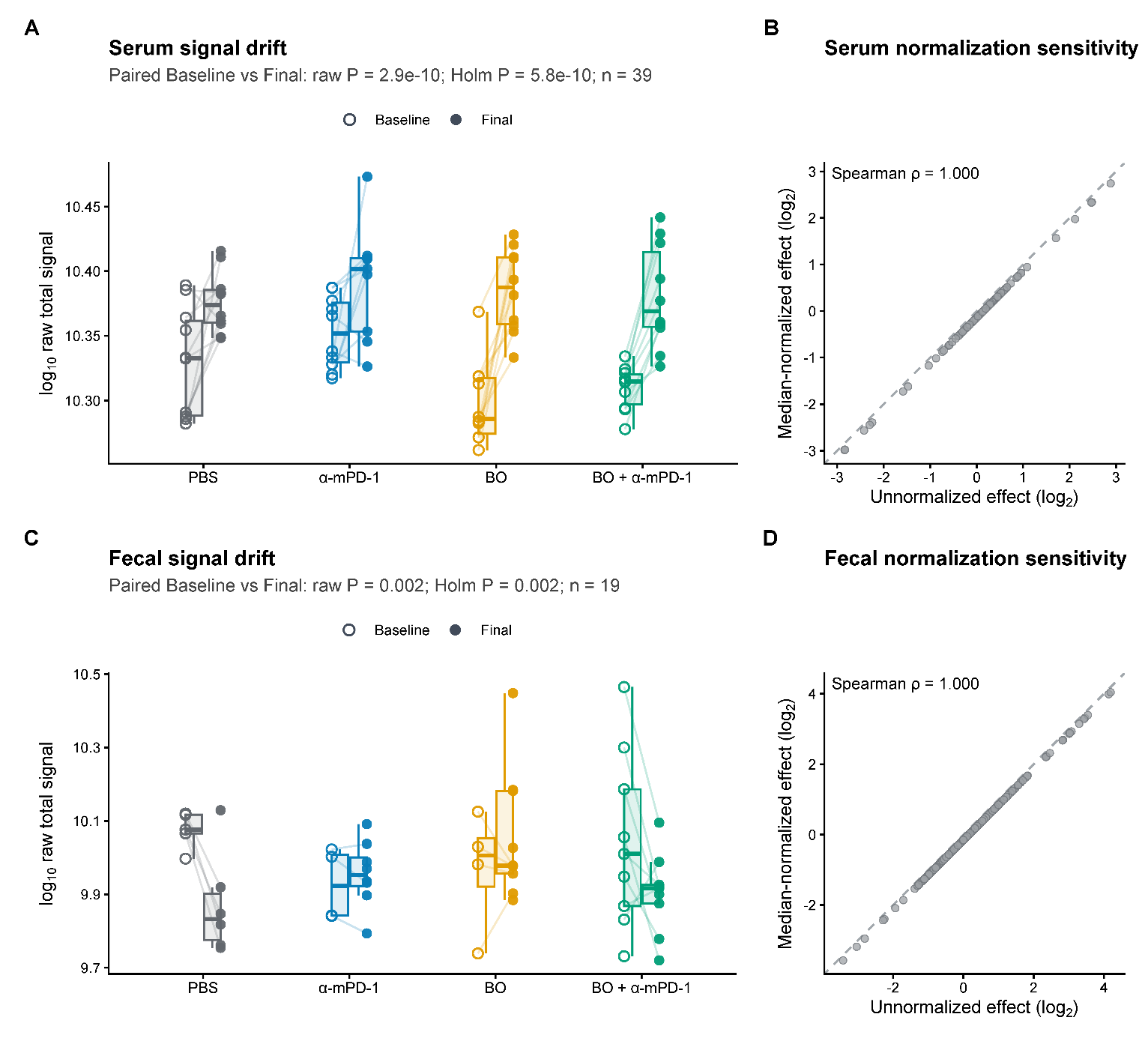
